# A single nuclei expression resource for exploring dog brain cell transcriptomic diversity

**DOI:** 10.64898/2026.09.14.751376

**Authors:** Matthew J. Christmas, Eric Pederson, Ola Wallerman, Paul Theodor Pyl, Susanne E. Reinsbach, Elisabeth Sundström, Chao Wang, Åsa Karlsson, Maja L. Arendt, Jennifer R. S. Meadows, Kerstin Lindblad-Toh

## Abstract

Domesticated dogs present a unique case in nature where mutualism with and selective breeding by humans has led to profound changes in their environment, physiology, and behaviour compared to their wolf-like ancestors. Dog tameness, sociability, and trainability have likely evolved due to significant alterations to brain function. Exploring the effects of domestication on the dog brain will be facilitated by the characterisation of dog brain cell diversity, a task which is not yet complete. To fill this gap, we use single-nucleus RNA sequencing and survey cell types across the dog brain. We present transcriptomes from almost 60,000 cells sampled from the cerebellum, thalamus, and three regions of the cerebrum. Our analysis identified 24 major clusters representing 21 broad cell types, and 131 subclusters revealing regional variation. Comparisons with human and mouse datasets revealed a high level of conservation in neurons across mammalian brains, and greater divergence in glial cells, particularly oligodendrocytes. We demonstrate the utility of the dataset for interrogating specific gene expression patterns across the brain, including genes implicated in dog domestication, behavioural traits, and disease. We discover distinct differences in the expression of opioid receptors in the brains of dogs, compared to humans and mice, providing a potential explanation for dogs’ higher tolerance and lower risk of severe adverse effects of opioid drugs. The data from this project are released via a web application for use by the wider community.

## Introduction

The mammalian brain is one of the most complex products of evolution, where the density and complexity of connections among a vast repertoire of neuronal and glial cell types results in advanced cognitive functions, including perception, coordination, problem solving, and consciousness^1,2^. Recent advances in single cell RNA sequencing methods^3^ are enabling the characterisation of the specific gene expression profiles across the catalogue of brain cell types. Comprehensive single cell atlases have now been developed for human^4^ and mouse^5^ brains, revealing the extreme diversity in cell types across brain regions as well as through developmental time^6^. These resources are providing novel insights into brain function^7,8^ as well as revealing cell types and molecular pathways involved in brain dysfunction^9–11^.

The domestic dog provides an interesting case for studying brain function. Its history of domestication from a wild wolf-like ancestor has likely had direct effects on brain function, with evidence for genetic alterations to neural crest and central nervous system development^12^. Signatures of parallel evolution in genes and gene pathways related to digestion^13^, metabolism, and neurological processes^14^ between dogs and humans have likely resulted from our shared history and environments. Selective breeding for specific traits in breed formation, such as herding and retrieving, has largely selected on ancient, non-coding regulatory variation, with much within-breed variation associated with neurodevelopmental co-expression networks^15^. Additionally, dogs can act as informative models of human brain disorders and dysfunction^16,17^ and for studying neurotoxicological side effects in the pharmacological industry^18^. Single cell resources are beginning to be developed for the dog: a single-cell RNA-seq atlas of healthy lung tissue from four dogs is now available^19^ as well as single-nuclei RNA-seq of hippocampus tissue from a single dog^20^. Development of further resources for additional brain regions is required to develop our understanding of the specific cellular lineages and molecular pathways that define canine brain architecture. Additionally, the degree of molecular homology between canine and human brain cell types is not yet established at a single-cell level, limiting the interpretability of the dog as a translational model for neurological research.

To fill this gap, we present the single-cell Dog Brain Expression resource (DBEx), a gene expression dataset encompassing over 59,000 cells from the cerebellum, thalamus and three regions of the cerebrum (frontal, occipital, and temporal lobes) from four beagle dogs, capturing the diversity of neuronal and glial cell types across these brain regions. We provide an overview of the resource and integrate it with similar datasets for human and mouse brains to explore similarities and differences in cell types and expression profiles across these mammals. We demonstrate the utility of DBEx for interpreting the functional relevance of outcomes from behavioural and disease-focussed genome-wide association studies (GWAS). We also consider the pharmaceutical relevance of the resource and explore expression profiles for drug targets, including opioid receptors. We have released the data from this project as a publicly available, interactive web application that can be queried by, for example, brain region, cell type, or gene with a user-friendly interface for exploring DBEx (https://dog-brain-single-cell.serve.scilifelab.se/app/dog-brain-single-cell).

## Results and discussion

### The single-cell dog brain expression resource - DBEx

Our dog brain single cell expression resource comprises 15 single-nuclei RNA-sequencing (snRNA-seq) libraries, generated using a combination of 10x Genomics and Parse Biosciences technology. Our snRNA-seq libraries represent five distinct brain regions (Fig. 1A; cerebellum and thalamus, and three regions of the cerebrum: frontal, occipital, and temporal lobes) from four beagle dogs, with at least two biological replicates (dogs) and three technical replicates (distinct slices) per region (Fig. S1; Table S1). The snRNA-seq data was processed through a stringent quality control pipeline to maximise the integrity of the data and ensure that the vast majority of retained cells were accurately assayed, reducing the impact of spurious signals from cell leakage and background expression (Tables S2-4; see methods). The decision to use two different sequencing technologies (10x and Parse) was made to maximise the amount of data we could generate within the constraints of the project. This comes with the tradeoff that technology-specific differences in the data generation can result in artifactual noise in the data that must be accounted for.

**Figure 1.**
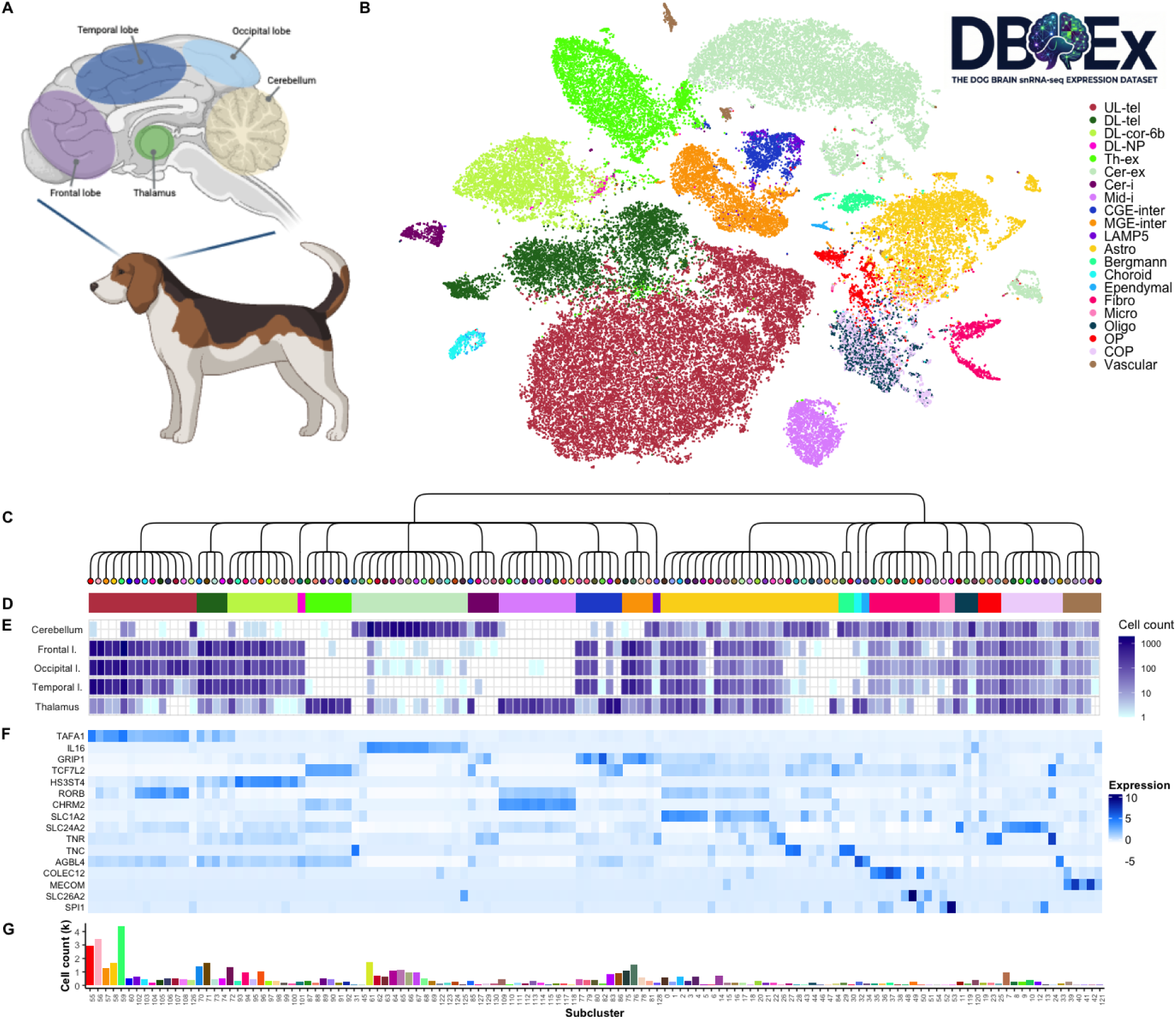
The dog brain expression resource. (A) Regions of the dog brain sampled for single nuclei RNA sequencing. (B) t-SNE plot showing clustering of harmonized expression data from 59,035 dog brain cells coloured by major cell types. Neurons: “UL-tel” = upper layer intratelencephalic, “DL-tel” = deep layer intratelencephalic, “DL-cor-6b” = deep layer corticothalamic and 6b, “DL-NP” = deep layer near-projecting, “Th-ex” = thalamic excitatory, “Cer-ex” = cerebellar excitatory, “Cer-i” = cerebellar inhibitory, “Mid-i” = midbrain-derived inhibitory, “CGE-inter” = interneurons from the caudal ganglionic eminence, “MGE-inter” = interneurons from the medial ganglionic eminence, “LAMP5” = LAMP5-LHX6 and Chandelier. Neuroglia: “Astro” = astrocytes, “Bergmann” = Bergmann glia, “Choroid” = choroid plexus, “Ependymal” = ependymal, “Fibro” = fibroblasts, “Micro” = microglia, “Oligo” = oligodendrocytes, “OP” = oligodendrocyte precursors, “COP” = committed oligodendrocyte precursors, “Vascular” = vascular. (C) Dendrogram representing the subclustering of cells into 74 neuron clusters across eleven major neuron cell types and 57 neuroglial clusters across ten major cell types. (D) Major cell types of subclusters are indicated by coloured bars, with colours matching the t-SNE plot in (B). (E) Cell counts per subcluster for each brain region. (F) Expression heat map showing expression of cell-type specific marker genes per subcluster. (G) Bar plot of total cell counts per subcluster, with the majority of cells sequenced in the study being neurons (81.3%). Logo in top-right of figure created using Google Gemini.

However, each of the five tissues contain data generated from both technologies, we saw similar cell-type clustering between samples from the same brain region regardless of technology, and we show high gene expression correlations between similar cells assayed using the two technologies (mean Pearson’s R = 0.90; Fig. S1).

### Assigning cell type labels

We assigned cell type labels to each cell in our dataset based on a correlation analysis, where gene expression profiles were compared to those of human cells^4^ using SingleR. We then took the informative Supercluster labels from the human data and assigned them to the dog cells based on the highest correlations (“cell types” from here on). Using this approach, 21 distinct cell types were identified across the five brain regions (Fig.1A,B). Spearman’s rank correlations between dog and human cells were generally higher for neurons than glial cells (mean maximum correlation = 0.56 and 0.40 for neuron and glial cells respectively; Fig. S2). This is as expected due to the high constraint of neurons among mammals^21^.

### Data harmonisation and clustering

We used Harmony^22^, implemented in Seurat v.4^23^, to harmonize the data across samples while accounting for non-biological signals relating to technical batch effects and performed clustering analysis. Cell clusters largely reflected the distinct cell types assigned by comparison to the human data. We used this cell assignment to filter out cells that were assigned cell type labels that did not match the label of the majority of cells in their neighbourhood, removing 6,279 cells. Whilst this approach results in information loss, it mostly removed sparse cells intermediate to major cell clusters in the tSNE plot. The final dataset contains transcriptomes for 59,035 cells (mean per-sample cell counts per technology: 10x = 5,067; Parse = 1,682), with 24 major clusters representing 21 broad cell types, and 131 subclusters revealing finer scale cell type distinctions (Figs. 1B-E, S3A,B, Table S5). Several clusters are limited to a single brain region, with the cerebellum and thalamus having region-specific cell clusters, while cortical regions generally share cell types (Fig. 1B-F).

### Neuronal cell types

The majority of sequenced cells were neurons (n = 48,016 cells, 81.3%) due to our enrichment of neuron nuclei in the library preparation (see Methods). Subclustering of all neuron cells revealed nine clusters, with cell type labels aligning well with the identified clusters (Fig. 2A; Fig. S3C). The clustering revealed region-specific neurons, with highly similar neuron types and proportions among the three sampled cortical lobes, as well as cerebellum- and thalamus-specific neurons (Fig. 1E). We assigned neurotransmitter identity to each of the main neuron cell clusters based on co-expression of canonical genes coding for neurotransmitter transporters and synthesising enzymes (Fig. 2B). The vast majority of neurons in our dataset are glutamatergic excitatory neurons (84.8%, n = 40,726), found across all five brain regions. The remaining 15.2% (n = 7,290) are GABAergic inhibitory neurons, including midbrain-derived inhibitory neurons found in the thalamus, cerebellar inhibitory neurons, and several classes of interneurons from the sampled cortical regions.

**Figure 2.**
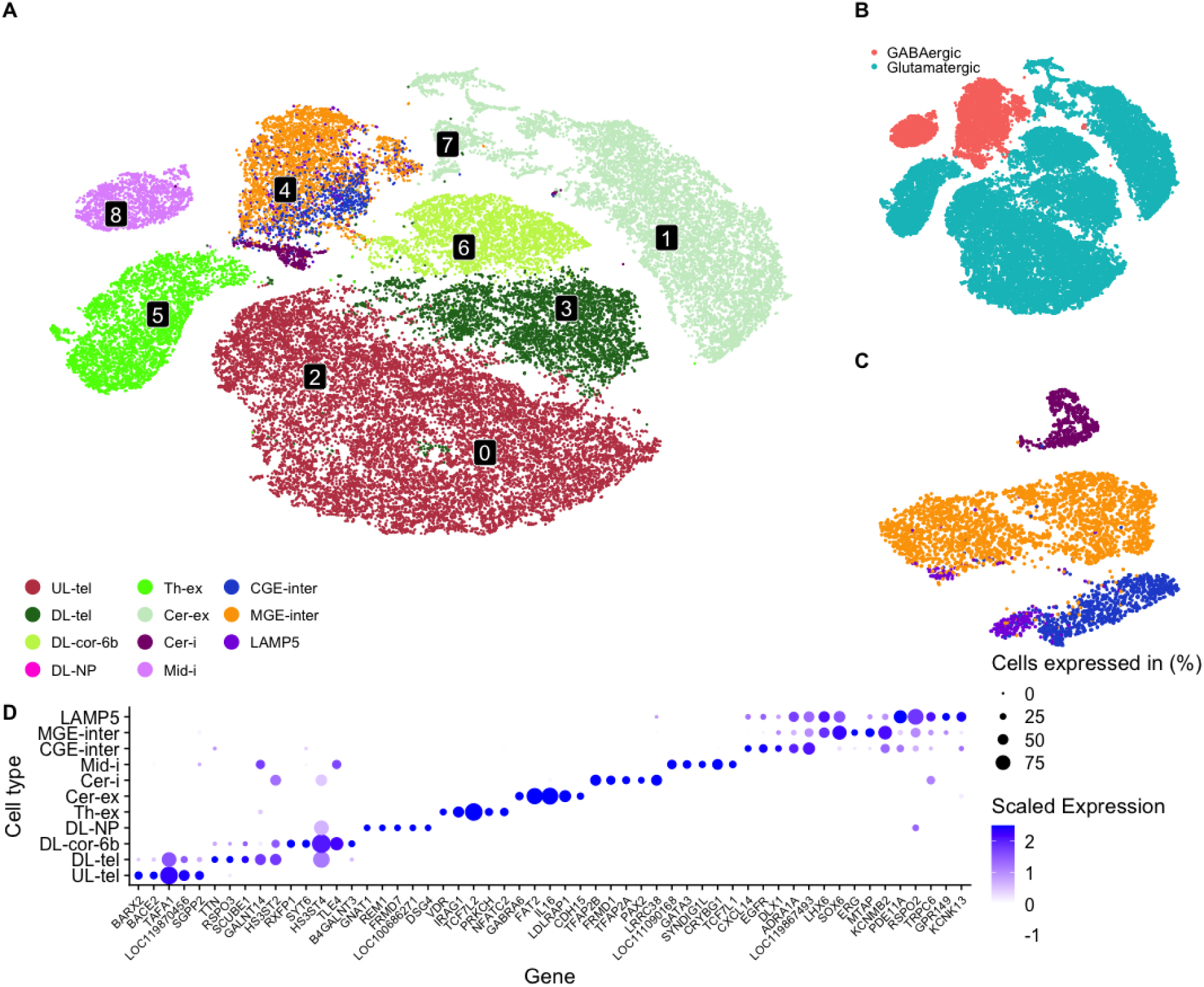
Neurons in the dog brain. (A) tSNE plot of neuron cells, coloured by cell type. Numbers indicate clusters identified using Harmony. B) tSNE plot with neurons coloured by their major neurotransmitter type. (C) tSNE plot showing subclustering of Harmony neuron cluster 4, containing GABA-ergic inhibitory neurons. (D) Dot plot showing expression of the top 5 differentially expressed genes per cell type, where genes were expressed in at least 25% of cells, showed a log2 fold change of > 0.5, and were upregulated in the focal cell type. Legend under (A) refers to colours in both (A) and (C). Cell type abbreviations are defined in the fig. 1 legend.

Neuron cluster four contained a mixture of inhibitory neuron types. Subclustering of these cells revealed three distinct populations of inhibitory neurons, one consisting of cerebellar inhibitory neurons, and two consisting of inhibitory interneurons predominantly from the cortex. This included a cluster of interneurons derived from the medial ganglionic eminence (MGE interneurons), and a cluster of interneurons derived from the caudal ganglionic eminence (CGE interneurons), as well as LAMP5-LHX6 and Chandelier interneurons (Fig. 2C). These cortical inhibitory interneurons share high expression of several genes that are not expressed in other neuron types, including the transcription factors *LHX6* and *SOX6*, which are both involved in interneuron development^24,25^ (Fig. 2D).

Thalamic neurons were distinct from those found in other regions and formed two clusters, one of excitatory (cluster 5) and one of inhibitory neurons (cluster 8); Thalamic excitatory neurons are signified by a high expression of *TFC7L2*, a transcription factor that is known to play a role in postmitotic development of glutamatergic neurons in the thalamus^26^.

Midbrain-derived inhibitory neurons from the thalamus showed high expression of *GATA3*, which is well characterised as a regulator of GABAergic neuron identity in the midbrain^27^. These cells also displayed specific co-expression of *CRYBG1, SYNDIG1L,* and *TFC7L1*, none of which has previously been identified as a canonical cell type marker and may represent dog-specific patterns. The vast majority of neurons from the cerebellum were excitatory granule neurons deriving from the upper rhombic lip (clusters 1 and 7), which show expression of the canonical granule neuron marker genes *FAT2* and *GABRA6.* Neuron cell types sampled from cortical regions included upper-layer intratelencephalic (clusters 0 and 2), deep-layer intratelencephalic (cluster 3), deep-layer near-projecting, and corticothalamic (cluster 6) neurons. Deep-layer neurons all show expression of *HS3ST4*, which encodes heparan sulfate 3-O-sulfotransferase 4, an enzyme that adds sulfate to heparan sulfate proteoglycans, key extracellular matrix components involved in maintaining neuronal excitability and plasticity^28^ (Fig. 2D).

### Non-neuronal cell types

Our dataset includes sampling of 11,019 glial cells (18.7% of all cells). Harmony analysis of all neuroglial cells resulted in the identification of 12 clusters, each broadly aligning with cell type labels: Astrocytes (clusters 0, 1, and 6), Bergmann glia (cluster 4), oligodendrocytes and committed oligodendrocytes precursor cells (cluster 2), oligodendrocytes precursors (cluster 3), ependymal cells (cluster 11), choroid plexus (cluster 7), microglia (cluster 10), fibroblasts (clusters 5 and 9), and vascular cells (cluster 8) (Fig. 3A, Fig. S3D). Astrocytes, oligodendrocytes and their precursor cells, vascular cells, and fibroblasts were found in all brain regions (Fig. 3B). Bergmann glia were found specifically in the cerebellum, where they function as highly specialised astrocytes involved in synaptic pruning, plasticity, metabolic support, and neuroprotection of Purkinje cells (reviewed in ^29^). Choroid plexus cells (specialized epithelial cells that produce and secrete cerebrospinal fluid) were detected in the thalamus samples. Close proximity of the choroid plexus and thalamus in the third ventricle likely explains the sampling of this cell type despite not targeting the choroid plexus in our sampling. Similarly, a small population of ependymal cells was also only sampled from the thalamus dissections, which likely included some of the surrounding ventricle where these cells are found. Microglia, the primary immune defense cells within the central nervous system, were only found in the occipital lobe dissections, which is not expected as this cell type is distributed widely throughout the brain and makes up ∼7% of glial cells in mammalian brains^30^. This anomaly is likely a result of the nuclei isolation method and enrichment for neurons. Significantly differentially expressed genes were identified for all cell types, supporting the clustering and labelling of cell types (Fig. 3C).

**Figure 3.**
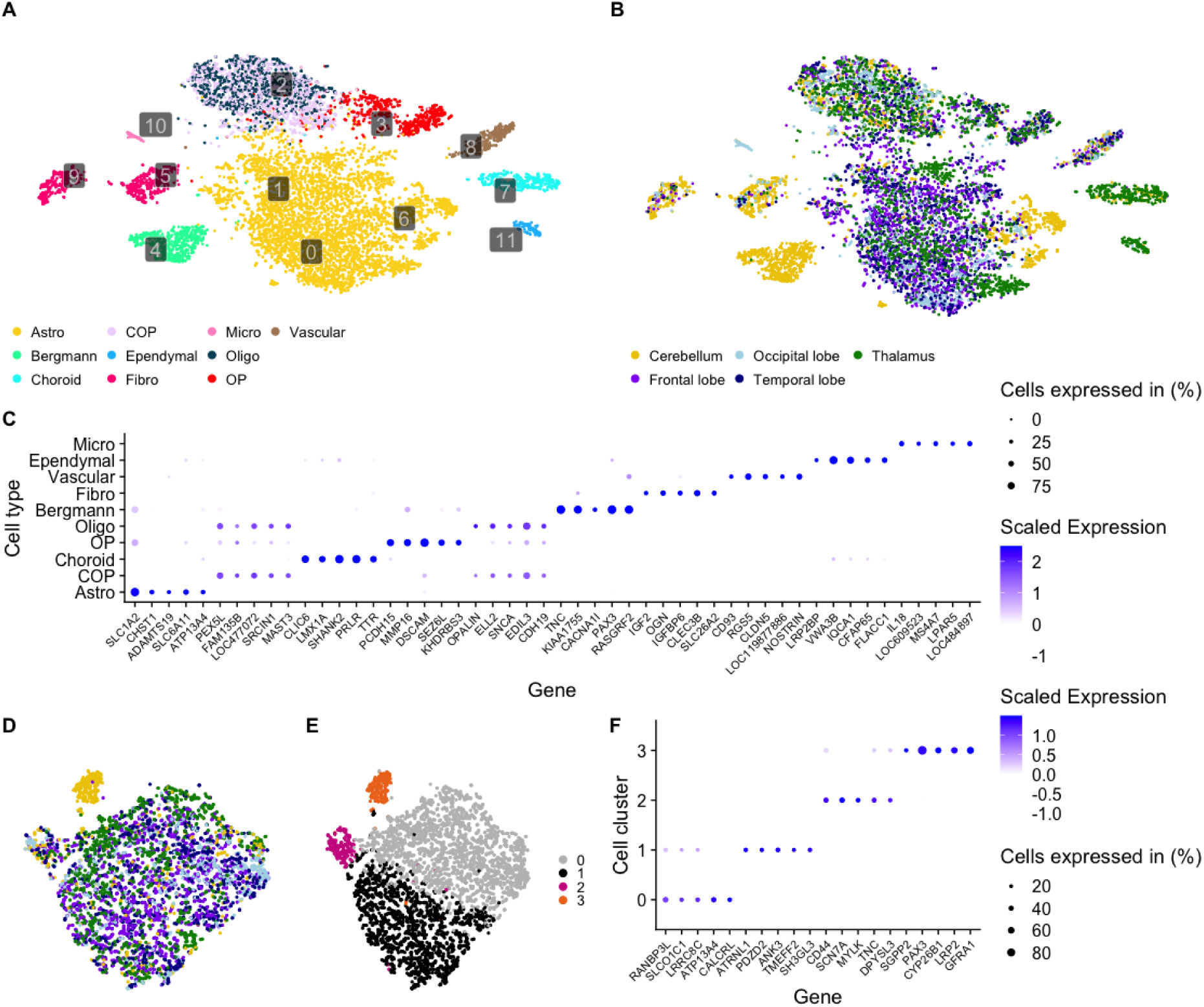
Neuroglial cells in the dog brain. tSNE projections of dog brain glial cells, coloured by (A) cell type and (B) brain region. Numbers in (A) indicate clusters identified using Harmony. (C) Dot plot showing expression of the top 5 differentially expressed genes per cell type, where genes were expressed in at least 25% of cells, showed a log2 fold change of > 0.5, and were upregulated in the focal cell type. tSNE projections showing subclustering of astrocytes coloured by (D) brain region and (E) identified Harmony clusters. Cluster 3 represents a population of cerebellum-specific astrocytes, distinct from cerebellar Bergmann Glia. (F) Dot plot showing expression of the top 5 differentially expressed genes per astrocyte cell cluster, where genes had to be expressed in at least 25% of cells, showed a log2 fold change of > 0.5, and were upregulated in the focal cell type. Cerebellum-specific astrocytes show high expression of *PAX3*, a gene encoding the PAX3 transcription factor that is highly expressed in Bergmann glial cells. Cell type abbreviations are defined in the fig. 1 legend.

Almost half of all sampled glial cells were astrocytes (42.9%, n = 4,728), which are a group of cell types involved in the maintenance of CNS homeostasis^31^. Re-clustering of only astrocytes revealed four subclusters (Fig. 3D,E). This included a small population of 205 cells specific to the cerebellum. Cells in this cluster showed high expression of *PAX3*, which is not expressed in astrocytes from other brain regions (Fig. 3F). The PAX3 transcription factor is a marker of Bergmann glia (Fig. 3C), a specialised astrocyte-like cell type specific to the cerebellum, which plays a critical role in supporting Purkinje neurons and thus located in the Purkinje layer^32,33^. In our data these 205 cells are distinct from Bergmann glia and likely represent a subclass of cerebellar astrocytes, potentially related to progenitor or specialised functional states distinct from Bergmann glia. Further investigation, including spatial transcriptomics, would be required to fully survey the repertoire of astrocyte-type cell populations in the dog cerebellum.

### Comparative expression profiles in other mammals

We integrated DBEx with single cell brain expression datasets from human^4^ and mouse^5^ to explore similarities and differences in gene expression across mammalian brains. We subsampled cells from the human and mouse datasets to produce equivalent datasets to ours, in terms of regions sampled and depth of sampling (see Methods). This resulted in a dataset of 59,035 cells per species (177,105 total), with mean gene counts per cell of 3,191 in dog, 2,269 in human, and 2,906 in mouse. Clustering of the integrated dataset resulted in the identification of 40 clusters (Fig. S4), with cell clusters largely containing equivalent cell types across the three species (Fig. 4A). Nearly all cell clusters contained cells from all three species (Fig. 4B), i.e. all cell types are represented in each species, except for one cluster containing almost exclusively mouse cells (labelled ‘mouse - mixed cluster’). However, there was clear species-specific subclustering of cells within certain clusters. To assess between-species cell type divergence further, we compared the average normalized expression values of orthologous genes and performed expression correlation analyses between cell types and species (Fig. 4C-F). Overall, dog and human cells had the highest average correlation (Pearson’s r = 0.74, 0.61, and 0.71 for dog-human, dog-mouse, and human-mouse). This is despite the greater divergence time between dogs and humans (∼94 mya) compared to humans and mice (∼87 mya; estimates from www.timetree.org), although it reflects the higher sequence similarity between dogs and humans compared to mouse and humans due to the faster evolutionary rate of mice. As expected due to their higher conservation across species, neurons showed higher correlations between species than glial cells (Pearson’s r = 0.81, 0.71, and 0.78 for neurons and 0.70, 0.52, and 0.61 for glial cells in dog-human, dog-mouse, and human-mouse comparisons respectively). Heatmaps of correlation values revealed generally higher correlations among the distinct neuron types, whereas glial cell types were generally more distinct from each other (Fig. 4C-F). The most divergent cell type between species based on these correlation scores was oligodendrocytes between dog and mouse (Pearson’s r = 0.39) and human and mouse (Pearson’s r = 0.53). Interestingly, dog-human oligodendrocytes showed among the highest correlations for glial cells (Pearson’s r = 0.80), suggesting higher similarity in this cell type between dogs and humans compared to mice.

**Figure 4.**
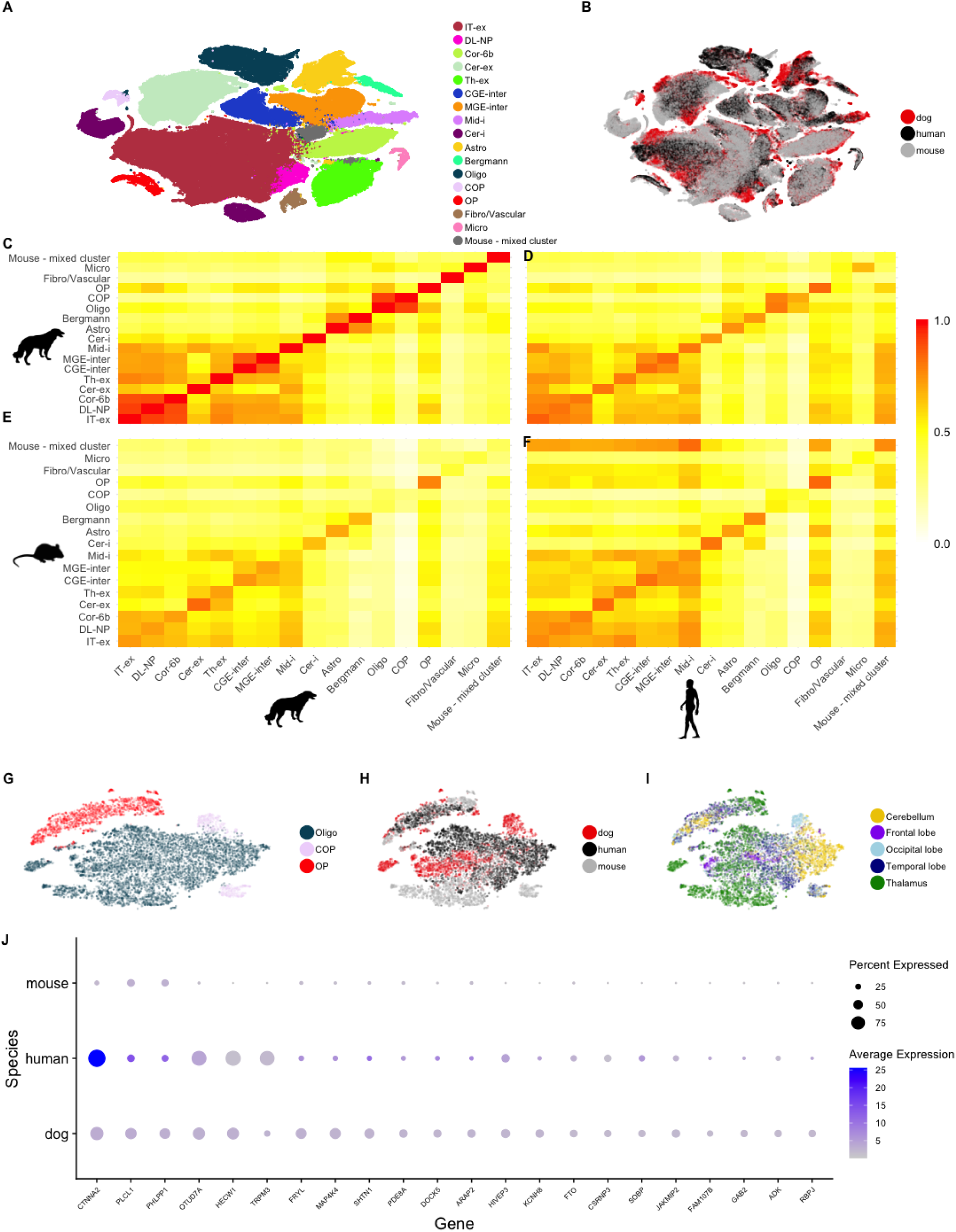
Comparative analysis of dog, human and mouse brain gene expression. tSNE plots showing integrated dog, human, and mouse single cell expression datasets coloured by (A) cell types and (B) species. Heat maps comparing gene expression between cell types in (C) dog versus dog, (D) dog versus human, (E) dog versus mouse, and (F) human versus mouse. tSNE plots of oligodendrocytes and their precursor cells from across the three data sets, coloured by (G) cell type, (H) species, and (I) brain region. (J) Dot plot showing expression of genes that are expressed significantly higher in dog and human oligodendrocytes compared to mouse oligodendrocytes. Cell type abbreviations are defined in the fig. 1 legend. Species silhouettes were obtained from phylopic.org.

Oligodendrocytes myelinate neuronal axons, which is essential for axon conductivity, as well as for supplying metabolites to neurons^34^. To investigate differences in expression between dog, human and mouse oligodendrocytes, we extracted all oligodendrocytes and their precursor cells (“OP”, “COP”) and reclustered them independently from other cell types (Fig. 4G). Oligodendrocyte precursor cells showed high similarity among all three species, with cells from all three species forming a single cluster (Fig. 4H; n = 571 from dog, 1,361 from human, and 791 from mouse). In each species comparison, oligodendrocyte precursors were consistently among the three most highly correlated cell types (0.84, 0.78, and 0.84 in dog-human, dog-mouse, and human-mouse comparisons respectively), demonstrating high conservation of this cell type. Differences in mature oligodendrocytes (n = 2,772 from dog, 5,671 from human, and 2,324 from mouse) were much more apparent in the clustering, with mouse cells showing little overlap with dog and human cells (Fig. 4H), as suggested by the correlation scores reported above. To investigate this further, we subsetted thalamic oligodendrocytes to focus on cells from a single brain region and reduce signals that may be due to differences across the brain rather than between species. The thalamus provided the greatest number of cells to compare (Fig. 4I; n = 1,219, 1,662, and 434 from dog, human, and mouse respectively). We then performed differential expression analysis to test for genes that were highly expressed in both dog and human oligodendrocytes but showed low expression in mouse oligodendrocytes. This analysis revealed 22 genes that showed significantly less expression in mouse cells compared to dog and human cells (Fig. 4J; log_2_ fold change < 0, p.adj < 0.05). These include fat mass and obesity-associated gene (*FTO)*, an eraser of m6A methylation on mRNA transcripts^35^. This RNA modification is highly enriched in the brain and is essential for oligodendrocyte maturation and axonal myelination^36^. The lack of *FTO* expression in mouse glial cells^37^ but presence in human and dog oligodendrocytes is suggestive of a more similar m6A demethylation mechanism in dog and human oligodendrocytes compared to mice. A variant in *FTO* has been associated with reduced brain size in humans, linking expression of this gene with brain development^38^.

The most significant differentially expressed gene was *CTNNA2/αN-catenin*, which encodes an adherens junction protein that plays a crucial role in the cadherin-catenin complex, where it links cadherins to the actin cytoskeleton predominantly in neurons^39^. The cadherin-catenin complex plays an important role in axon-oligodendrocyte contact and myelination in mammals^40^, a process essential to neurological function. Loss of *CTNNA2* causes structural abnormalities in human brains, including pachygyria (unusually thick cerebral folds/gyri)^41^, which may relate to its comparatively low expression in mice, which are lissencephalic and lack gyri^42^. Mutations in and around the gene have been associated with multiple human psychiatric disorders and phenotypes, including Alzheimer’s disease^43,44^, schizophrenia^45^, major depressive disorder^46^, negative thinking and obsessive compulsive disorder (OCD)^47^, impulsivity^48^, and excitement seeking^49^. Intriguingly, it has also been implicated as a candidate gene in a canine model of OCD^50^.

Differences in gene expression between human and mouse oligodendrocytes, including that of *CTNNA2*, have previously been identified^51^,supporting the results we present here. The accelerated evolution that appears to have driven the divergence of human oligodendrocytes from those in our primate cousins^52^ means that human oligodendrocytes are less conserved than other brain cell types. Our results of high correlation between human and dog oligodendrocyte expression profiles suggest that dogs may therefore present as better models than mice for human brain function and dysfunction, especially where it relates to oligodendrocyte function, although wider sampling of dog breeds is required to confirm this. The similar expression of a gene that has been highly implicated in overlapping human and dog brain disorders calls for further exploration of the role played by the CTNNA2 protein in oligodendrocyte-neuron interactions in these species.

### Species specific expression of opioid receptors

Single cell RNA sequencing methods are being utilised for drug development, helping to identify drug targets and aiding preclinical disease model choice, as well as delivering new insights into drug mechanisms of action^53^. In this context, we used DBEx to explore the expression of certain drug receptors in the dog brain to identify expression locality in comparison with expression in human and mouse brains. In particular, we examined the expression of the four types of opioid receptor family genes - *μ* (*OPRM1)*, *κ* (*OPRK1*), *δ* (*OPRD1*), and nociceptin receptor (*OPRL1*) - in our integrated dog-human-mouse dataset. This revealed a stark contrast in the dog brain compared to human and mouse: the dominant receptor in human and mouse brains is the *μ*-receptor, with high expression of *OPRM1* in multiple neuron types, especially cerebellar and thalamic excitatory neurons. However, the dog brain shows relatively low expression of the *μ*-receptor gene (Fig. 5A-E). In contrast, the dominant receptor in the dog brains we sampled is the *δ*-receptor, with high expression of *OPRD1* in cerebellar excitatory neurons, whereas this gene shows almost no expression in human and mouse cerebellar excitatory neurons (Fig. 5F-H). Opioids are commonly used as analgesics in both humans and dogs, with most clinical opioids including morphine, fentanyl, tramadol and codeine, targeting *μ*-opioid receptors^54,55^. However, the effective dose of opioid drugs is significantly higher in dogs and some drug formulations have limited effect. For example, the analgesic efficacy of tramadol in dogs has recently been shown to be low and potentially ineffective^56,57^. Lethal doses of opioids can be orders of magnitude higher in dogs compared to humans: LD_50_ of intravenous fentanyl is 0.03 mg/kg in primates, whereas it is over 450x higher in dogs at 14 mg/kg^58^. Similarly, the lethal dose of intravenous morphine for an adult human is estimated to be 30-50 mg total (∼0.4-0.7 mg/kg for an 70 kg adult)^59^ whilst for dogs the LD_50_ it is estimated to be 133mg/kg^60^. In humans, opioid overdose leads to respiratory failure due to *μ-*receptor binding in the brainstem leading to depression of rhythmic breathing^61^. Although we do not specifically target the brainstem here, our observed overall lower expression of *OPRM1* and higher expression of *OPRD1* in our dog samples compared to humans and mice can provide an explanation for the much lower efficacy and higher tolerability of opioid drugs in dogs. If these opioid expression patterns are common among dogs, which requires confirmation, then this would suggest that dogs do not represent valid comparative preclinical models for studying the pharmacologic effects of opioids intended for human use and that opioid analgesics that target *δ* opioid receptors may be more effective in dogs, which should be investigated further^62^.

**Figure 5.**
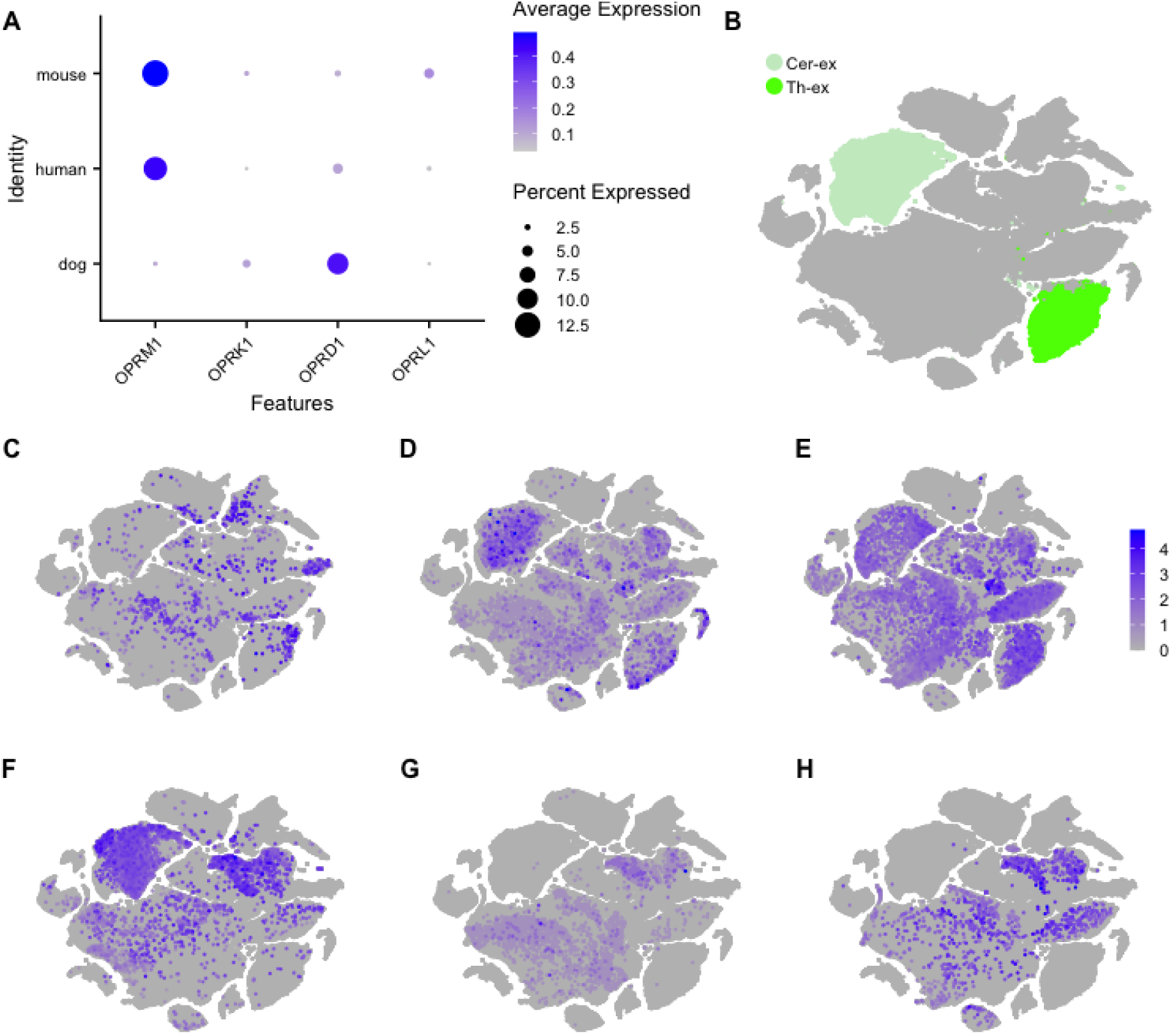
Expression of opioid receptor genes in dog, human and mouse brain cells. (A) Dot plot showing expression of opioid receptor genes across the three species data sets. (B) tSNE of integrated cells as in Fig. 5A, used here to highlight the locations of cerebellar excitatory and thalamic excitatory neurons, where the three species mainly differ in their expression of opioid receptor genes. (C-H) tSNE plots of all integrated cells showing *OPRM1* expression in (C) dog, (D) human, and (E) mouse cells, and *OPRD1* expression in (F) dog, (G) human, and (H) mouse cells.

### Expression of genes associated with dog domestication

Previous studies have presented evidence for selection on loci that altered expression of genes involved in CNS development during dog domestication^12,13,63^. Our dataset presents an excellent resource for exploring where these domestication-related genes are expressed in the adult dog brain. We queried our dataset with the 215 genes that are located within 50 Kbp of 58 candidate domestication regions (CDRs) identified from comparisons between village dog and wolf genomes^12^. Out of these 215 genes, 160 (74%) are detected as expressed in the dog brain, with 123 (57%) meeting a conservative threshold of expression in at least 10% of cells of at least one cell type (Fig. 6a).

**Figure 6.**
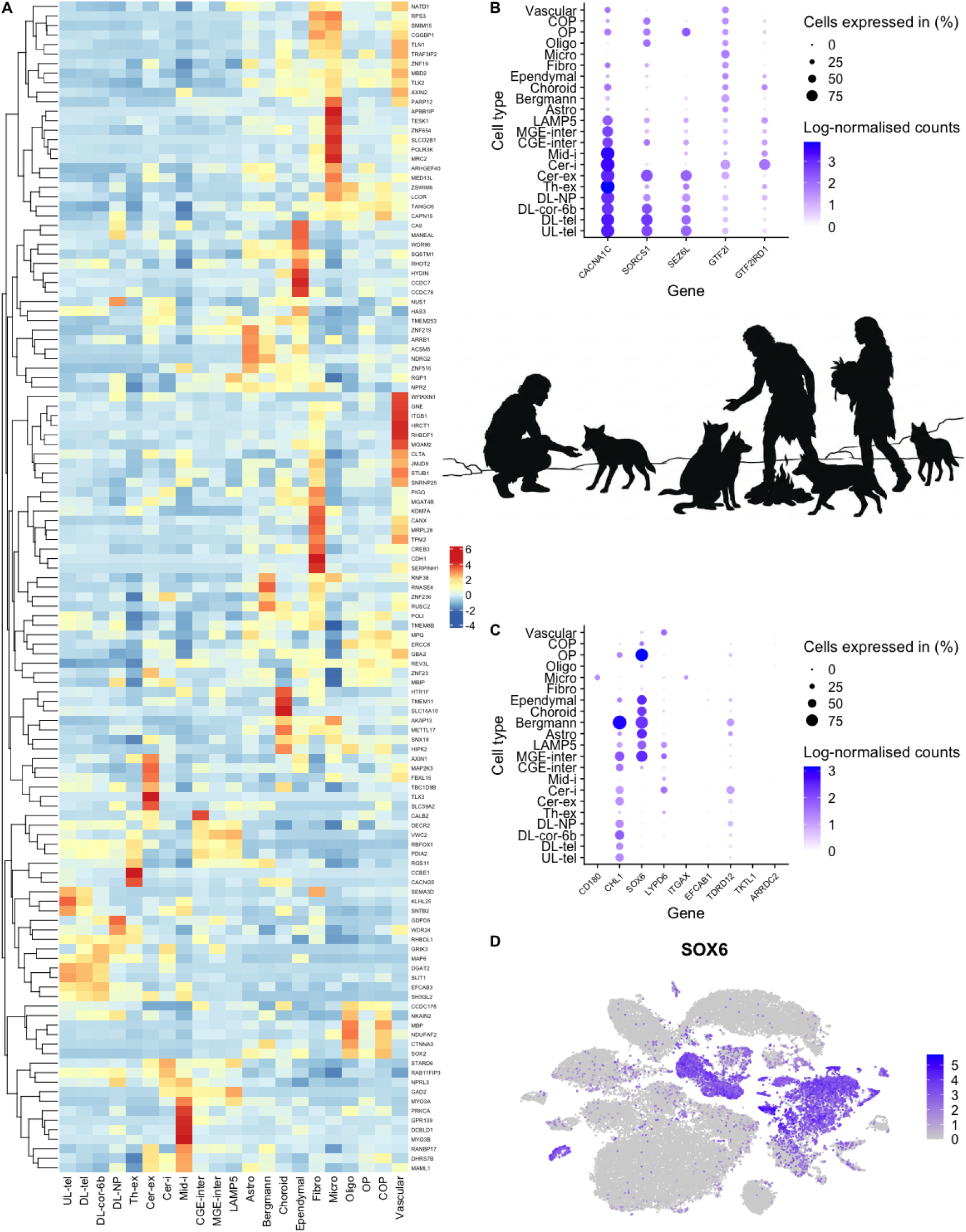
Expression of domestication-related genes across the dog brain. (A) Heat map of per cell type relative expression of 123 genes within 50 Kb of Candidate Domestication Regions (CDRs) and are expressed in at least 10% of cells of at least one cell type. Dendrogram on the y-axis shows hierarchical clustering of genes based on their expression patterns across cells. (B) Per-cell expression levels of genes associated with tameness and sociability in dogs. (C) Expression of genes shown to be differentially expressed between dog and wolf frontal cortex. (D) Expression of *SOX6* across all cells in the dataset. Cell type abbreviations are defined in the fig. 1 legend. Image panel generated using Google Gemini.

Our dataset reveals groups of CDR-associated genes that demonstrate cell-type specific expression, suggesting that the functions of specific cell types have evolved during dog domestication (Fig. 6a). For example, several genes show high expression in excitatory neurons in the cortex, including *SLIT1*, which is known to be involved in axonal growth^64^, and *SEMA3D*, which encodes the axon guidance molecule Semaphorin 3D^65^. Interestingly, missense mutations in *SEMA3D* and the related *SEMA3E* have previously been identified as behaviour-altering mutations during the domestication of pigs^66^. *RBFOX1* also shows high expression in dog neurons, except for cerebellar and midbrain-derived inhibitory neurons.

This gene encodes the RNA binding protein Fox-1 Homolog 1 and its expression has been associated with levels of aggression in several species^67^. Several genes, including *MYO3A* and *MYO3B*, *PRKCA*, and *DCBLD1*, show specifically high expression in midbrain-derived inhibitory neurons (Fig.6A). These neurons, sampled from the thalamus in our dataset, play key roles in controlling sensory processing, motor control, and decision making. The anatomical position of the thalamus in the diencephalon is key to its role in integrating signals from many areas of the CNS and influencing whole-brain activity and adaptive behaviour^68^.

We also observe CDR-related gene sets that show glial cell-type specific expression, particularly in microglia, with a cluster of seven CDR-related genes showing microglial-specific expression (Fig.6A). Microglia are the immune cells of the brain and their development has been shown to be sensitive to environmental changes, including alterations to the microbiome and maternal immune activation^69^. They display a conserved core gene program across mammals, but in humans there is significant heterogeneity and disruption to their function is implicated in several degenerative disorders, including Alzheimer’s and Parkinson’s disease^70^. In our SingleR analysis, the lowest correlations were observed between dog and human microglia, suggesting high divergence of the gene expression pathways (Fig. S2C). Unfortunately, the limited sampling of microglia in our dataset limits further analysis here and the fact that this cell type is highly context sensitive calls for much wider sampling of microglia to demonstrate true species-wide expression patterns. But the fact that we see high expression of multiple CDR-related genes in the four dogs we sampled calls for further investigation of this cell type in future studies.

Increased tameness and human sociability during domestication has been associated with several genes, in dogs^71^ and other species, including *CACNA1C* (dog sociability^72^)*, SORCS1* (tameness in foxes^73^)*, GTF2I* and *GTF2IRD1* (dog hypersociability^74^), and *SEZ6L* (dog sociability^75^). We see high expression of these genes across multiple cell types in the brain (Fig. 6B). In particular, *CACNA1C* shows high expression in all neuron types. This gene encodes the Cav1.2 subunit of L-type Ca2+ channels and as such plays a crucial role in calcium-mediated processes in neurons. It is one of the most highly replicable susceptibility genes for human neuropsychiatric disorders, including impaired social and cognitive processing^76^. Previous comparisons of bulk RNA expression data from dog and wolf brains have revealed multiple differentially expressed genes (DEGs)^63,77^. In particular, Albert *et al*., (2012)^77^ identified 30 DEGs between dog and wolf frontal cortex, 19 of which are upregulated in dogs. Interrogation of our DBEx resource revealed the cell types these genes are expressed in (Fig.6C), such as *SOX6* expression in cortical interneurons, glial cells (OPCs and Astrocytes), and in non-neural barrier cells of the choroid plexus and ependyma (Fig.6D). *SOX6* is a regulator of cell fate and differentiation in the CNS, controlling the timing of cell maturation, especially for oligodendrocytes and interneurons^78,79^. Increased *SOX6* expression in dog brain cells may contribute to the neoteny (the retention of juvenile physical or behavioral traits in an adult organism) seen in domestic dogs, as part of the domestication syndrome^80^. For example, in OPCs, *SOX6* represses terminal differentiation into mature, myelinating oligodendrocytes^81,82^. Similarly, in MGE-derived interneurons, inhibitory cells that regulate the synchrony and rhythm of cortical circuits, *SOX6* is crucial for their normal positioning and maturation^78^. The previously demonstrated upregulation of *SOX6* in dog brains compared to wolves in combination with our results here, showing high expression in diverse cell types, points to a systemic regulatory change in the dog genome that may affect brain developmental timing across multiple cell lineages. This presents an interesting avenue for future research to test whether increased *SOX6* expression in dogs leads to prolonged or delayed maturation of these cells, potentially contributing to differences in cognitive processing speed or development observed between dogs and wolves and ultimately supporting the suite of behavioral and physical traits that distinguish dogs from wolves, including reduced fear, increased social flexibility, and other behavioral traits characteristic of dog tameness^77^.

### Exploring expression of canine brain disorder related genes

We extracted gene names for all genes associated with dog nervous system diseases in the Online Mendelian Inheritance in Animals (OMIA) database (www.omia.org; accessed 01.11.2025), giving a list of 87 genes (Table S6). We then performed per-cell-type bulk expression analysis to identify the cell types in which each gene is expressed in DBEx (Fig. 7A). The strongest cell type-specific expression we observe is in microglia, where there is high microglia-specific expression of several OMIA genes implicated in neurodegenerative disorders, all of which involve degraded states of axonal myelin, including degenerative myelopathy (*SP110*), polyneuropathy (*SBF2*), hypomyelination (*FNIP2*), neuroaxonal dystrophy (*RNF170, VPS11*), *PCYT2* deficiency (*PCYT2*) and neuronal ceroid lipofuscinosis (*TPP1*, *CLN5*). Microglial action has been linked to neurological dysfunction and de-myelination in dogs, including in canine degenerative myelopathy^83^ and distemper^84^. It has recently been shown in humans and mice that microglia are essential for the regulation of myelin growth and preservation of myelin integrity by preventing its degeneration in adulthood^85^. Altered function of microglia, the brain’s primary immune cells, will therefore likely disrupt the integral role they play in myelin preservation, leading to neurodegenerative disorders. As we have previously discussed, we unfortunately only achieved a small sample of microglia cells and so these findings require confirmation from further sampling across multiple dogs of different breeds. Despite this, other recent findings have revealed the importance of microglia to canine cognitive dysfunction and they are emerging as key therapeutic targets for disorders involving myelin degradation in dogs^86^.

**Fig 7.**
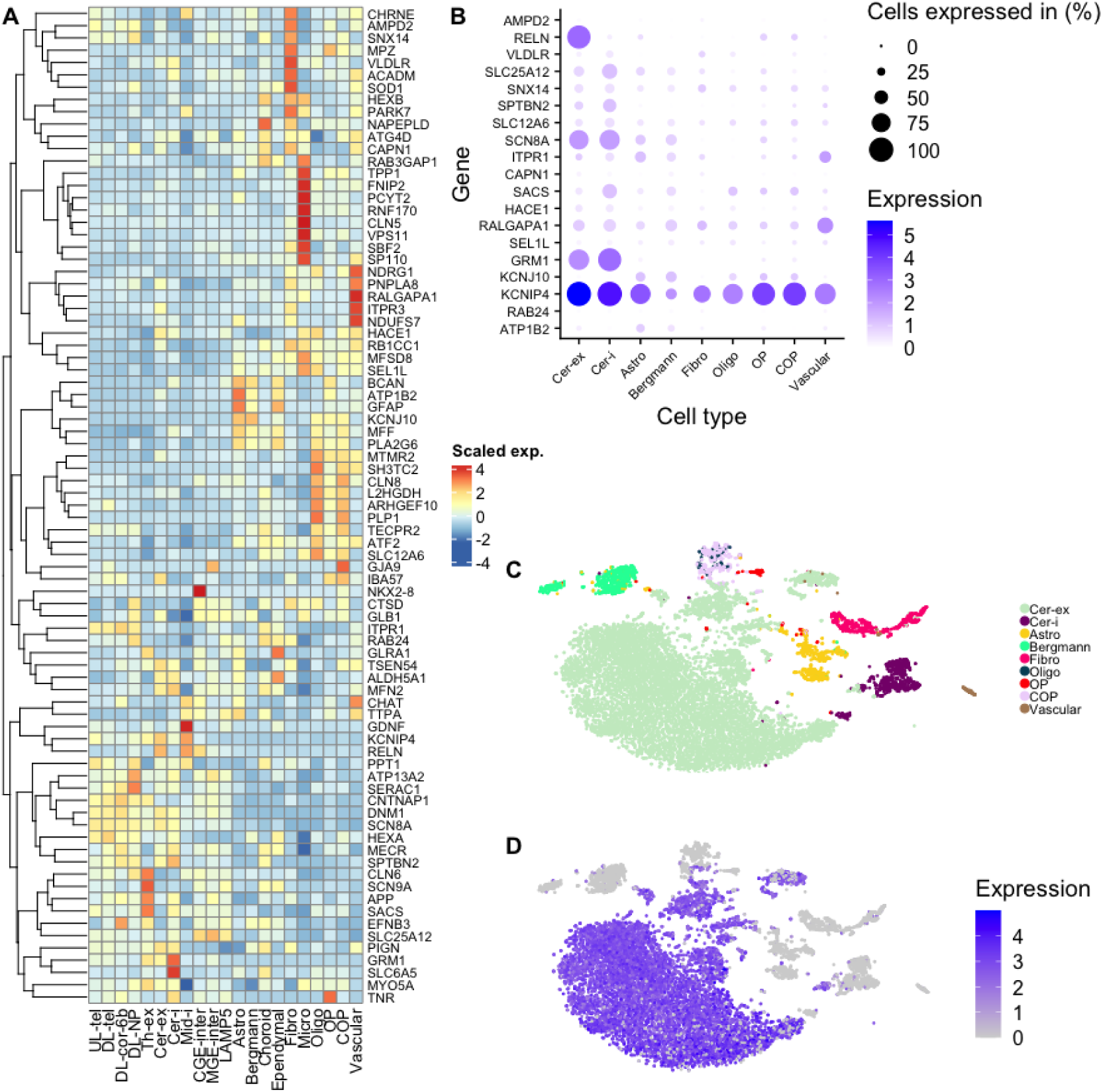
Expression of neurological disease-implicated genes across brain cell types. (A) Per cell type relative expression of genes implicated in nervous system-related phenotypes in dogs in the Online Mendelian Inheritance in Animals (OMIA) database. Hierarchical tree on Y-axis indicates expression profile similarity among genes (B) Expression of genes implicated in disorders of the cerebellum across cell types in the dog cerebellum. (C) tSNE plot of dog cerebellum cells. (D) High expression of *RELN* is seen specifically in cerebellar excitatory neurons in the cerebellum. The gene encodes a large secreted extracellular matrix protein involved in control of cell-cell interactions critical for cell positioning and neuronal migration during brain development. A frameshift deletion in this gene causes cerebellar hypoplasia in white swiss shepherd dogs. Cell type abbreviations are defined in the fig. 1 legend.

We next extracted all genes associated with canine disorders of the cerebellum from OMIA, giving a list of 21 genes, 15 of which are associated with cerebellar ataxia (Table S7). Disorders of the cerebellum generally lead to reduced motor coordination and imbalance in dogs. We see expression of 19 cerebellum disorder-associated genes in DBEx, most of which are broadly expressed across multiple cell types (Fig. 7B). The gene *RELN*, which encodes the protein reelin, showed the most specific cell type expression, with expression observed in the majority of sampled cerebellar granule cells and low or no expression in all other cell types (Fig.7C,D). Reelin is involved in controlling cell-cell interactions and is critical for cell positioning and neuronal migration during brain development^87^. A frameshift deletion in this gene has been associated with lissencephaly and cerebellar hypoplasia in white Swiss shepherd dogs, leading to progressive ataxia from 2 weeks of age^88^. Autopsies of affected brains revealed disorganised cerebellums, with thin and irregular granular layers^88^. Although DBEx does not capture developmental stages, the specific expression of *RELN* revealed here suggests that reelin continues to play an important role within mature granule cells of the cerebellum and alterations to its expression may lead to granule cell dysfunction and disease. In humans, abnormal expression of reelin is also associated with ataxia^89^, as well as multiple neuropsychiatric disorders, including autism, schizophrenia, bipolar disorder, major depression, and Alzheimer’s disease^90^. We see similar locality of *RELN* expression across dog, human and mouse brains (Fig. S5), suggesting a conserved function for this gene.

### Cell-type expression of dog behaviour-related genes for developing functional hypotheses

There have been several recent efforts to identify loci associated with various behavioural traits in dogs (e.g.^15,91,92^). Here, we demonstrate the utility of DBEx for exploring expression of trait-associated genes discovered through GWAS, providing supporting evidence towards the role specific genes may play based on the locality of their expression. Morrill *et al.,* (2022)^91^ identified 11 genome-wide significantly associated variants to behavioural traits. The most significantly associated locus was in a region containing the gene *SNX29*, which codes for a sorting nexin protein involved in intracellular trafficking and signaling. This region was associated with the phenotype “gets stuck behind objects”. We see broad expression of this gene across all cell types, with especially high expression in midbrain-derived inhibitory neurons from the thalamus (Fig. 8A). In rodents and humans, thalamic dysfunctions are linked to perseveration, navigational deficits, and disorders of movement initiation^93,94^. Testing whether dogs that display this ‘stuck’ behaviour show alterations in expression of *SNX29* in brain cells, and particularly mindbrain-derived inhibitory neurons, would be the next step in deciphering the cause of this phenotype.

**Figure 8.**
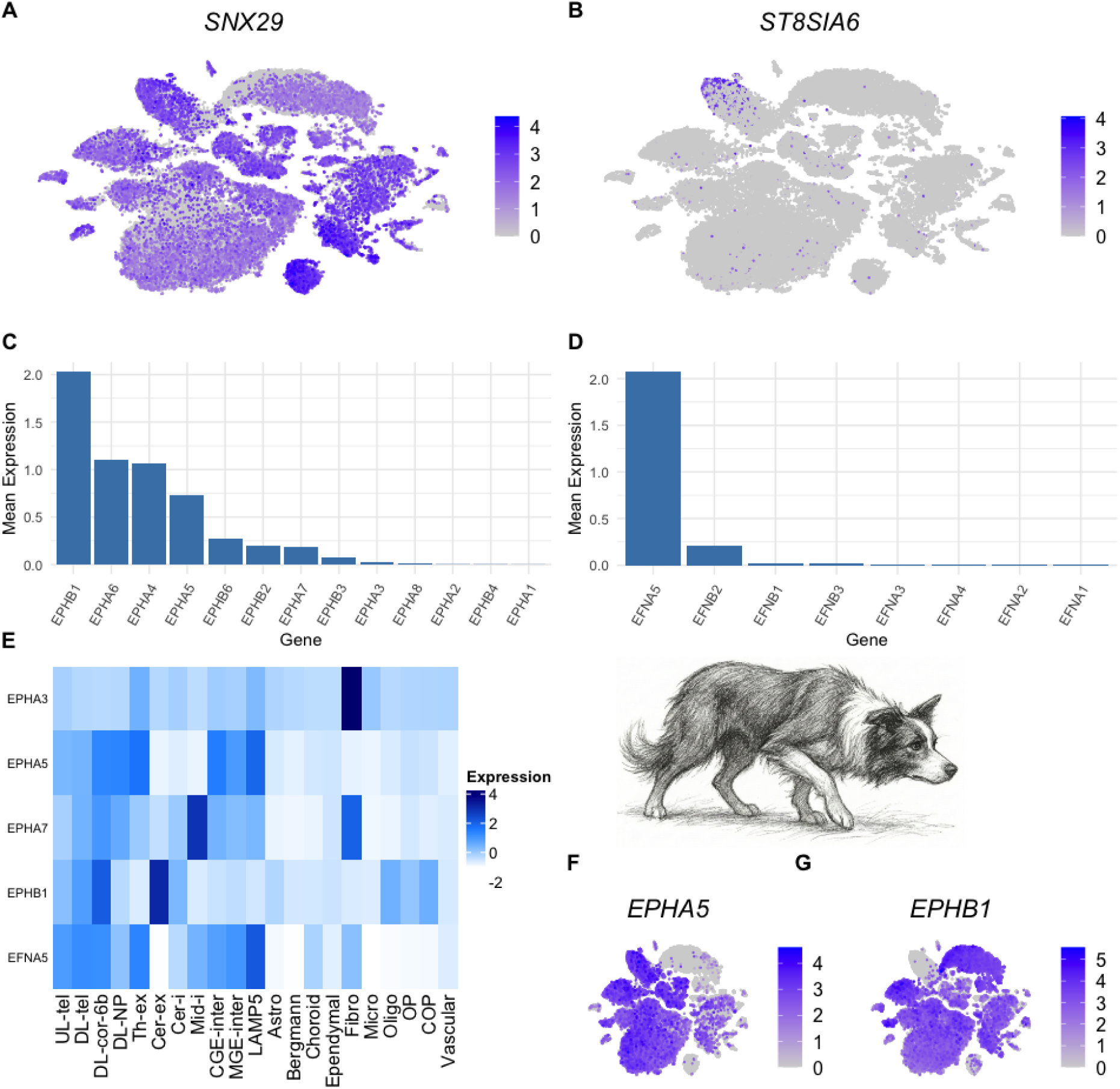
Expression of herding behaviour associated genes. (A) Cell-specific expression of *SNX29*, a gene associated with dogs becoming stuck behind objects. (B) Expression of *ST8SIA6*, variation in which is associated with sociability towards humans. Expression of this gene is mostly localised to a subset of thalamic excitatory neurons. Mean expression of (C) EPH receptor genes and (D) their ligands across all cells in DBEx. (E) Relative expression per cell type of ephrin genes associated with herding behaviour. (F) Expression of *EPHA5* is seen in most neuron types except those in the cerebellum and midbrain-derived inhibitory neurons. (G) *EPHB1* shows particularly high expression in excitatory neurons of the cerebellum, a brain region involved in object tracking and movement. Cell type abbreviations are defined in the fig. 1 legend. Image panel generated using Google Gemini.

The same study also found a significant association between a region within the second intron of *ST8SIA6* and the level of sociability towards humans. This gene codes for a sialic acid transferase, which adds a2,8-linked disialic acids on O-linked glycoproteins (sialylation). These marks on glycoproteins can then bind to the inhibitory receptors on innate immune cells, suppressing immune activation^95^. The role of this specific gene in neurons is unknown, but levels of sialylation are high in the brain and have been associated with several neurological disorders in humans (reviewed in^96^). We see specific, high expression of *ST8SIA6* in three subclusters of thalamic excitatory neurons (clusters 88, 89 and 91), with very little expression observed in any other cell type (Figs. 8B; S6). Thalamic function is involved in social behaviour in mammals, with a prefrontal-thalamic circuit associated with sociability in mice^97,98^, and the number of thalamic neurons is strongly associated with ASD-type behaviour, including lower sociability, in rats^99^. Further exploration of the role of ST8SIA6 in thalamic excitatory neurons is required, but it may be acting to signal to microglia, which carry out autophagy or ‘synaptic pruning’ to refine brain circuits and do not destroy cells carrying these disialic acid marks^100^. Indeed, it has been shown that disruptions to microglia-mediated synaptic pruning impairs functional brain connectivity and social behaviour^101,102^.

Variation in ephrin receptor genes (*EPH*) and the genes encoding ephrin ligands (*EFN*) is associated with herding behaviour in dogs, including receptor genes *EPHA3, EPHA5, EPHA7, EPHB1,* and ligand gene *EFNA5*^15,92^. Ephrin receptors are transmembrane receptor tyrosine kinases (RTKs) that bind membrane-bound ephrin ligands expressed by neighboring cells, mediating short-range cell-to-cell communication and play multiple roles in neural development^103,104^. Ephrin signalling has been demonstrated to trigger apoptosis, regulating the number of neural progenitors and ultimately altering brain size and shape^105^. We calculated mean expression (counts per cell) of all ephrin receptor and ligand genes to identify which genes are highly expressed in DBEx (Fig. 8C,D). The most highly expressed *EPH* and *EFN* genes were *EPHB1* and *EFNA5*, both of which are among the genes associated with herding behaviour and both show high expression across multiple cell types and regions (Fig. 8E-G). Haplotypes of *EPHA5* and its ligand *EFNA5* have previously been associated with herding behaviour in sheepdogs^15^. We see co-expression of these two genes across all neuron types except for neurons of the cerebellum and midbrain-derived inhibitory neurons (Fig. S7).

*EPHB1* shows broad expression across multiple cell types, with high expression in the excitatory neurons in the cerebellum (Fig. 8E,G). Haplotypes within this gene show segregation within breed lineages for border collie show dogs versus working dogs and a working line-specific haplotype is associated with increased levels of chase-bite motor patterns^92^. Ephrin-B1 is present in rat granule cells during early post-natal development and its overexpression enhances survival and neurite growth in cultured cerebellar granule neurons^106^. The cerebellum plays a key role in the control and coordination of movements, as well as the tracking of moving objects^107^, and so cerebellar neurons likely play important roles in shaping herding behaviour in dogs, potentially involving the expression of *EPHB1* during development. Our demonstration of neuron-specific expression of ephrin receptor genes and their ligands provides further support to the hypothesis that these genes are involved in shaping dog herding behaviour through their regulation of neuronal growth, warranting further investigation.

### Limitations

There are several limitations to the dataset we present here, not least the sampling of only four dogs from the same breed. Whilst this potentially reduces the generalisability of some of our results, the dogs were all healthy at the time of sampling and, as we demonstrate, single cell expression profiles were highly consistent among samples, as well in comparison to human and mouse cells. We therefore suggest that, whilst there are bound to be individual differences in gene expression among samples, overall cell-type specific expression can be retrieved from this limited sampling and is likely highly representative of dogs more widely. However, caution must be used when considering differences in expression of specific genes without further confirmatory evidence, such as in our opioid receptor example. We also used two different snRNA-seq approaches to generate the data, which may have led to technical biases. However, QC and integration of the datasets resulted in limited technological bias and we demonstrate high consistency among similar cell types generated from the two approaches, again providing high confidence in the accuracy and robustness of the dataset.

## Conclusions

The Dog Brain Expression resource provides a high resolution profile of single-nuclei gene expression in neuronal and glial cells across the dog cerebral cortex, thalamus, and cerebellum. We have demonstrated the utility of this resource for exploring the expression of genes related to domestication across the dog brain. The contrasting patterns of opioid receptor gene expression we find in our dog samples compared to human and mice samples provides a potential explanation for the great differences in sensitivity to opioid drugs between these species. Despite this difference, our comparative analysis of dog, human and mouse brains demonstrates a higher similarity between the expression profiles of dog and human cells overall compared to mice, particularly in oligodendrocytes. If wider sampling of more dogs across more breeds confirms our findings then the domestic dog may emerge as a more suitable model over mice for preclinical drug trials, and dogs have the potential to provide pharmacological meaningful trials with lower attrition rates and better prediction of potential side effects.

## Methods

### Sampling

Four dogs, two male and two female beagles, were sampled in this study (Table S1). They were sourced from a commercial provider, Zyagen, USA. All dogs were housed in a breeder facility which meets the USDA Animal Welfare Act standards and American Kennel Club standards. Dogs were all healthy at the time of sampling, were housed in spacious and ventilated conditions with opportunities for exercise and socialisation, and received a consistent feeding regime with high quality nutritional food as well as regular health checks and vaccinations. We therefore do not expect that the environmental conditions of the dogs will have had significant impacts on the gene expression analyses we present in this study. Dogs were euthanised prior to sampling by a licensed veterinarian using a sedative followed by injection of sodium pentobarbital.

### Tissue & Extraction

Extracted brains from the four dogs were frozen whole at the time of extraction. Brain tissue was later partially thawed and dissected by a veterinarian and neuroscientist with 40 years experience in *in vivo* research, utilising standardised dog brain atlases for accurate anatomical reference. Dissected sections were then stored at −80°C.

### Single Nuclei RNA-Seq

Nuclei extraction was performed using 2 ml Dounce homogenizer (KIMBLE) and pestle method where pestle ‘A’ performed 10 strokes and pestle ‘B’ performed 20 strokes. The resulting solution of nuclei and nuclei extraction solution was then filtered through a 30 μm filter (Miltenyi). The homogenization buffer for nuclei isolation was prepared fresh with 10 mM Tris pH 8.0, 250 mM sucrose, 25 mM KCl, 5mM MgCl2, 0.1% Triton-X 100, 0.5% RNasin Plus RNase inhibitor (Promega), Halt Protease Inhibitor Cocktail (100X; ThermoFisher Scientific) and 0.1 mM DTT. After centrifuging the nuclei to produce a pellet (400 xG at 4°C), the nuclei were resuspended in the antibody buffer containing 1X PBS, 0.8% nuclease-free BSA (Bovine Albumin Fraction V (7.5% solution, ThermoFisher Scientific)) and 0.5% RNasin Plus RNase inhibitor. The NeuN Antibody, anti-human/mouse/rat, APC, REAfinity (Miltenyi Biotec, 130-119-493) was added to the nuclei and antibody buffer solution at 10 ul for every 10^7^ nuclei and samples were incubated for 30 minutes at 4°C. After primary antibody incubation, samples were centrifuged for 5 minutes at 400 xG and 4°C to pellet nuclei and pellets were resuspended in 1X PBS with 0.8% BSA and 0.5% RNasin Plus. Next, the secondary antibody (goat anti-mouse IgG (H+L), Alexa Fluor 594 conjugated, ThermoFisher Scientific, A-11005) was applied to nuclei suspensions at a dilution of 1:2.5 for 30 minutes at 4°C. Nuclei suspensions were then centrifuged at 400 xG and 4°C for 5 minutes. An aliquot of 20 μL of Anti-PE MicroBeads UltraPure (Miltenyi Biotec) per 10^7^ total nuclei was then added with 80 μL of LS buffer (degassed PBS, pH 7.2, 0.5% BSA, and 2 mM EDTA) and subsequently centrifuged for 5 minutes at 400 xG and 4°C to pellet nuclei. The pellet was resuspended with 500 ul of LS buffer (10^8^ nuclei per 500 μl LS buffer) before applying the sample to a washed LS column (Miltenyi Biotec) with 3 ml placed in the MidiMACS Separator (Miltenyi Biotec). Subsequent washing of the sample in the LS column with 3 ml of LS buffer 3 times was followed by the immediate flushing of the sample by removing it from the magnetic separator and applying the plunger into the column. Afterwards the number of nuclei were counted and evaluated using a Bürker chamber (BLAUBRAND) and trypan blue stain (Invitrogen) before either; (i) directly handing in the samples to the National Genomics Infrastructure (NGI) facility to be processed with the Chromium Single Cell 3’ Reagent Kit (Version 3, 10X Genomics), (ii) performing the steps in the Chromium Single Cell 3’ Reagent Kit (Version 3, 10X Genomics) in house, (iii) applying 200 – 500 μl of freezing medium (70% glycerol in buffer N; 10 mM Hepes pH7.5, 2 mM MgCl2, 25mM KCl, 250 mM sucrose, containing 1 mM DTT, 1 mM PMSF, 1X Halt Protease Inhibitor Cocktail) and storing the sample in the −80°C freezer or (iv) running the Evercode Fixation kit (Parse Biosciences). The nuclei suspension mostly had high quality nuclei, with some signs of debris or lower quality nuclei in some cases. For extracted nuclei that were frozen in the buffer, they were thawed at 37°C for 5 minutes before being diluted with PBS and 0.1% BSA and centrifuging (400 xG at 4°C), decanting and resuspending with PBS and 0.1% BSA. Sorted and unsorted samples were then combined in a 9:1 ratio (sorted:unsorted) to enrich neuronal nuclei so that the samples were not dominated by glial cells^4^.

### 10X

Single-cell RNA sequencing was performed using the Chromium Single Cell 3’ Reagent Kit (Version 3, 10x Genomics) following the manufacturer’s protocol. Briefly, single-cell suspensions were loaded onto the Chromium Controller to create Gel Bead-in-Emulsions (GEMs) for barcoding individual nuclei. Reverse transcription and cDNA amplification were carried out prior to library construction. Paired read libraries with an insert size of 150 bp were prepared according to the manufacturer’s instructions for sequencing on the Illumina NovaSeq 6000.

### Parse Biosciences

Single-cell RNA sequencing was also performed using the Evercode Whole Transcriptome kit (Parse Biosciences) according to the manufacturer’s protocol. Briefly, single-cell suspensions were fixed and permeabilized prior to three rounds of split-pool combinatorial barcoding. Following barcoding, libraries were constructed using both barcoded random and oligo-dT primers to enable broad transcript coverage. Paired read libraries with an insert size of 150 bp were prepared according to the manufacturer’s instructions for sequencing on the Illumina NovaSeq 6000.

### Data pre-processing and QC

Output sequencing FASTQ files from the 10X protocol were processed using cellranger count v.7.1.0^108^, with the ‘include introns’ setting, and the Parse data were processed with splitpipe v. 1.1.0, defining the chemistry and kit used for each library. These pipelines perform alignment, filtering, barcode counting, and UMI counting. Raw reads were aligned to the augmented reference UU_Cfam_GSD_1.0-Y from the Dog10K project^109^ with gene annotation information from the NCBI gene annotation 106 (available at https://ftp-ncbi-nlm-nih-gov.ezproxy.its.uu.se/genomes/all/annotation_releases/9615/106/). Output count matrices from both the 10X and Parse pipelines were then processed through several quality control steps, detailed below, with the aim of retaining only reliable, high fidelity cells.

### Removing cells with excess spliced transcripts

We ran Velocyto^110^ on each samples’ data to calculate the number of unspliced transcripts per cell. As we used snRNA-seq, the majority of transcripts should be unspliced so this provides a useful filter to remove cells where the majority of mRNA has not come from the nucleus, indicated by the presence of spliced transcripts. We therefore removed any cells with a proportion of unspliced transcripts of less than 50%.

### Removing likely ambient/background RNA

We used the remove-background tool in Cell-Bender v.0.3.2^111^ to identify and remove likely background RNA from the counts matrices. We ran remove-background in GPU mode (flag ‘cuda’) and used default settings for the false positive rate (0.01), low count threshold (5), and epochs (150). We adjusted the learning rate in each case based on the interpretation of the learning curve, as suggested in the guide (https://cellbender.readthedocs.io/en/stable/usage/index.html). After background removal, Cellbender outputs were converted to Seurat objects for further quality control in R.

### Further quality filtering

Counts in each Seurat object were log normalised using the Seurat tool NormalizeData (method = “LogNormalize”). We tested for sample mixup by checking that only males showed expression of genes on the Y chromosome. We filtered out cells with a number of features (genes) below 100 or above median plus two times the median absolute deviation. We used DoubletFinder in R to identify and remove doublets, where two or more cells share the same cell-identifying barcode. This method is sensitive to identifying ‘heterotypic doublets’, which are doublets formed from transcriptionally-distinct cell states, but is unlikely to detect homotypic doublets (doublets formed from transcriptionally-similar cell states). Following the QC filter, all Seurat objects were merged into a single Seurat object for downstream analysis, containing expression data for 21,241 genes.

### Cell type labelling

We used SingleR (https://bioconductor.org/packages/release/bioc/html/SingleR.html) to label cell types in our data based on the similarity between their expression profiles and the expression profiles of cells in the human cell atlas^4^. SingleR calculates Spearman rank correlations between the single-nucleus profiles and the human reference dataset. To ensure high-confidence assignments, SingleR performs an iterative fine-tuning process and prunes labels where the ‘delta’ (the difference between the top-scoring label and the median of all labels) is significantly low by iteratively removing labels with scores more than 0.05 below the maximum, effectively requiring the best match to be distinct from the background distribution of scores.

### Harmonization and clustering

A principal component analysis (PCA) was run on the log-normalised expression values of the final set of cells using the RunPCA function in Seurat. We used the ElbowPlot function to create a scree plot and assess how many PCs likely retain biological information and should be retained in downstream analysis, retaining 50 PCs. The data were then harmonized using the RunHarmony function, grouping by sample ID for batch correction (<u>group.by</u>.vars=”sample_id”) and by technology (10x and Parse). RunUMAP and RunTSNE were then run on 30 dimensions using the harmony reduction (dims = 1:30, reduction = “harmony”)., followed by FindNeighbours on the harmony reduction, giving K-nearest neighbour (KNN) and shared nearest neighbour (SNN) graphs. We used the SNN graph as input to FindClusters for cell clustering. This same approach was used for all clustering of subsets of the data (e.g. clustering of neuronal and neuroglial cells separately). To determine the optimal clustering resolution, we utilized an iterative approach. Multiple resolutions were tested and evaluated by visualizing the resulting clusters on UMAP embeddings and assessing the expression of known canonical marker genes. The final resolution for each subset was chosen to ensure the segregation of biologically distinct cell populations while avoiding the over-clustering and fragmentation of transcriptionally homogeneous states (see Table S8 for the full list of dimensions and resolutions used).

### Neurotransmitter types

We examined the expression of canonical neurotransmitter transporter genes and synthesising enzymes to assign neurotransmitter types to neurons. Neurotransmitter genes queried were: *SLC17A6, SLC17A7, and SLC17A8* for glutamertergic (Glut), *SLC32A1, SLC18A2, GAD1, GAD2,* and *ALDH1A1* for GABAergic (GABA), *SLC6A5* for glycinergic (Glyc), *SLC18A3* and *CHAT* for cholinergic (Chol), *SLC6A3, SLC18A2, TH*, and *DDC* for dopaminergic (Dopa), *SLC6A4, SLC18A2*, *TPH2*, and *DDC* for serotoninergic (Sero), *SLC6A2*, *SLC18A2,* and *DBH* for noradrenergic (Nora), and *SLC18A2* and *HDC* for histaminergic (Hist). We checked for coexpression of both neurotransmitter transporter and corresponding transmitter synthesising enzymes, requiring an expression value of log_2_(counts per million) > 3 to assign neurotransmitter identity to each neuron cluster. In our dataset, we only identified GABAergic and Glutaminergic neurons, which is expected given the cell types we sampled.

### Generating the multi-species comparative dataset

We integrated our dog data with snRNA-seq data from the most comprehensive human brain expression atlas^4^ and single cell RNA-seq data from the most comprehensive mouse brain atlas^5^. We subsampled the human and mouse datasets in R to match the brain regions, cell counts, and mean UMI counts achieved in the dog dataset to increase comparability. Mouse gene IDs were converted to human IDs to match the other two datasets by extracting mouse-human orthologs from Ensembl Biomart and then replacing mouse gene IDs with human gene IDs for all orthologs in the Seurat object in R, removing all rows for genes without orthologs. We filtered the gene sets to retain only 1:1 orthologs between all three species using ortholog status defined for each gene in^112^. We then normalized and scaled each of the three datasets using NormalizeData and ScaleData in Seurat v.4 and ran FindVariableFeatures with features=2000. We identified genes common to all three datasets using Reduce in R and then ran prepSCTIntegration with the common genes list as the anchor.features. We ran PCAs on each dataset using the common genes using RunPCA and then ran FindIntegrationAnchors using “SCT” as the normalization method, “rpca” as the reduction, and common genes as the anchor features to create the integrated dataset for downstream analysis. We then ran Harmony, UMAP, tSNE, FindNeighbours, and FindClusters on the integrated dataset as described for the dog dataset.

### Cross-species cell type expression correlations

To compare cell-type expression profiles across species, we first extracted per-species datasets from the integrated Seurat object and calculated bulk average gene expression per cell type using AverageExpression on the SCT-transformed data. For each species comparison, we generated correlation matrices in R and calculated Pearson’s correlation coefficient of gene expression per cell-type. Cell type correlations were then plotted as heat maps using ggplot2.

### Oligodendrocytes analysis

All cells labelled as oligodendrocytes, COPs, or OPs were subsetted from the integrated dataset and clustered using FindNeighbours with the first 10 PCs from the PCA and FindClusters with a resolution of 0.2. We extracted all mature oligodendrocytes from the thalamus and then tested for DEGs using FindMarkers, with mouse cells set as ident.1 and dog and human cells set as ident.2, using a Wilcoxon signed rank test and setting ‘min.pct = 0.1, logfc.threshold = 0.25’. We then filtered the DEGs for genes that show similar average expression between dogs and humans. We ran AverageExpression on the SCT assay grouped by species and then filtered for genes where the absolute expression difference between human and dog cells was < 0.2 and the difference in mouse was > 0.5 and then filtered for only those genes significantly upregulated in dog and human cells (average log2FC < 0, adjusted p value < 0.05).

### Statistical analysis and plotting

All statistical analyses were performed in R v.4.4.1 “Race for your life” using the packages Seurat v4, tidyverse, and ggplot2 for plotting. tSNE plots of cell clustering were generated using DimPlot in Seurat. We used a combination of the inbuilt analysis and plotting tools in Seurat, including DotPlot, FeaturePlot and VlnPlot, to explore cell type-, region-, and species-specific expression levels of genes previously implicated in dog domestication, behaviour, and disease, as well as genes with pharmacological relevance (i.e. opioid receptor genes).

### Data web application

We have made the DBEx data available via an online application created using the R package ShinyCell2^113^. This package creates a Shiny app from the Seurat dataset in R, which is then hosted by SciLifeLab Serve. This allows users to visualise the dataset and query specific genes to explore their expression patterns in the dog brain.

## Data availability

Raw snRNA-seq reads are available from the European Nucleotide Archive (ENA) via accession number PRJEB108828. The processed dataset is available for exploration via a web app: https://dog-brain-single-cell.serve.scilifelab.se/app/dog-brain-single-cell.

## Code availability

Code for filtering and analysing the snRNA-seq data is available on GitHub at https://github.com/MattChristmas/DBEx.

## Acknowledgements

We thank Sten Linnarsson and Kimberly Siletti for useful discussions during the planning and data analysis stages of the project respectively. Computation and data handling were enabled by resources provided by the Swedish National Infrastructure for Computing (SNIC) at Uppsala Multidisciplinary Center for Advanced Computational Science (UPPMAX projects, UPPMAX2025/2-118 and UPPMAX2025/2-119), partially funded by the Swedish Research Council through grant agreement no. 2018-05973. Some laboratory work was performed using the National Genomics Infrastructure Sweden (NGI). Bioinformatics support was provided by the National Bioinformatics Infrastructure Sweden (NBIS). Technical infrastructure for hosting the DBEx app was provided by SciLifeLab Serve (https://serve.scilifelab.se), a platform developed and supported by SciLifeLab Data Centre.

## Author contributions

Project conception: K.L.T

Laboratory work: E.P., O.W., E.S.

Bioinformatics: P.T.P, S.R, C.W, M.J.C

Analyses: M.J.C

Data interpretation: M.J.C, J.R.S.M, M.L.A, E.P, K.L.T

M.J.C wrote the first draft and all authors edited the manuscript.

