## Supplementary figures for "A single nuclei expression resource for exploring dog brain cell transcriptomic diversity"

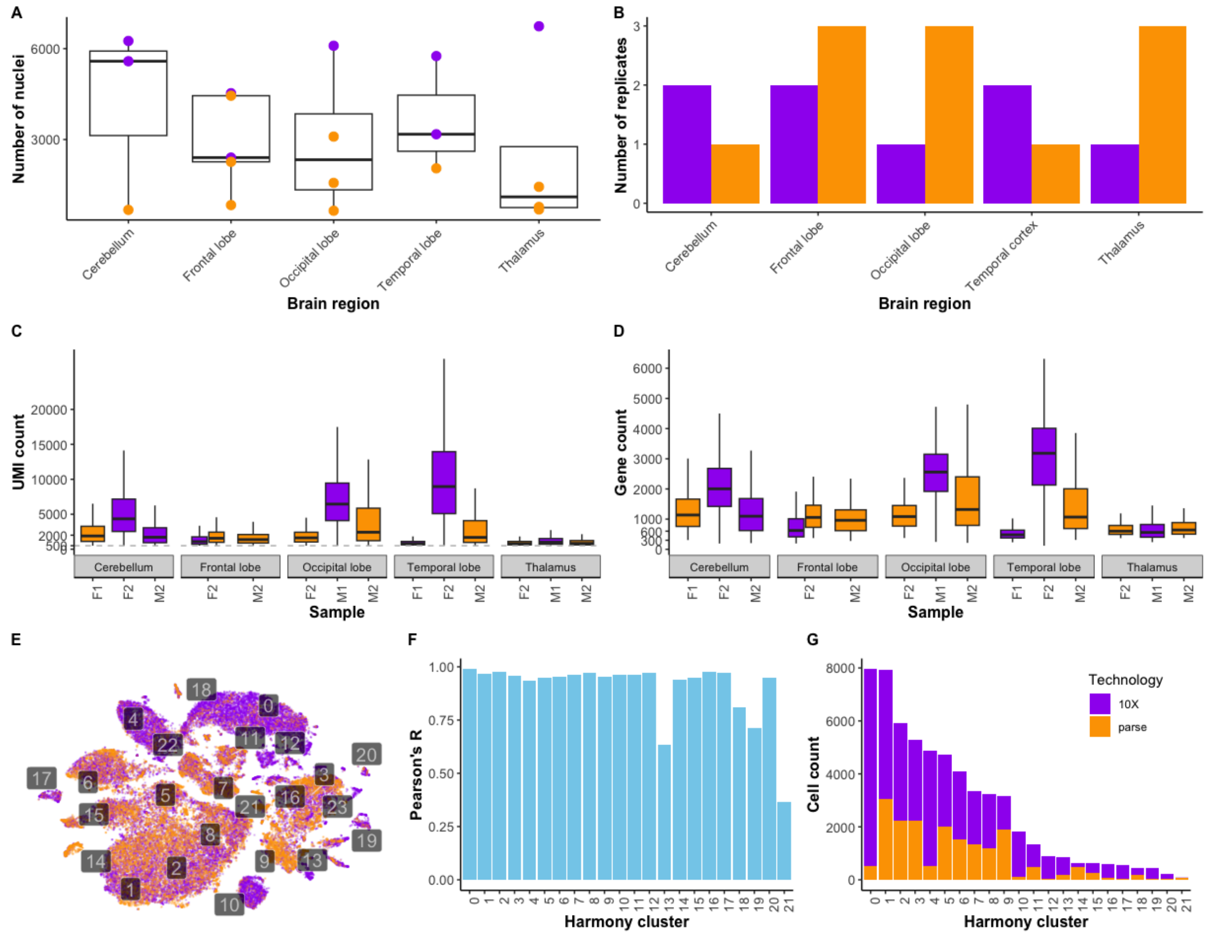


**Figure S1. Overview of sampling and data.** (A) Boxplots of the number of nuclei sampled per brain region per technology. (B) Bar plots showing the number of technical replicates per brain region and technology. Boxplots showing (C) UMI counts and (D) gene counts per sample per brain region, coloured by technology. (E) tSNE plot of integrated data set, with cells coloured by technology. Numbers indicate Harmony clusters. (F) Bar plot showing the Pearson's correlation between average gene expression of cells generated using each technology, per Harmony cluster. (G) Stacked bar plot of cell counts per Harmony cluster, coloured by technology.

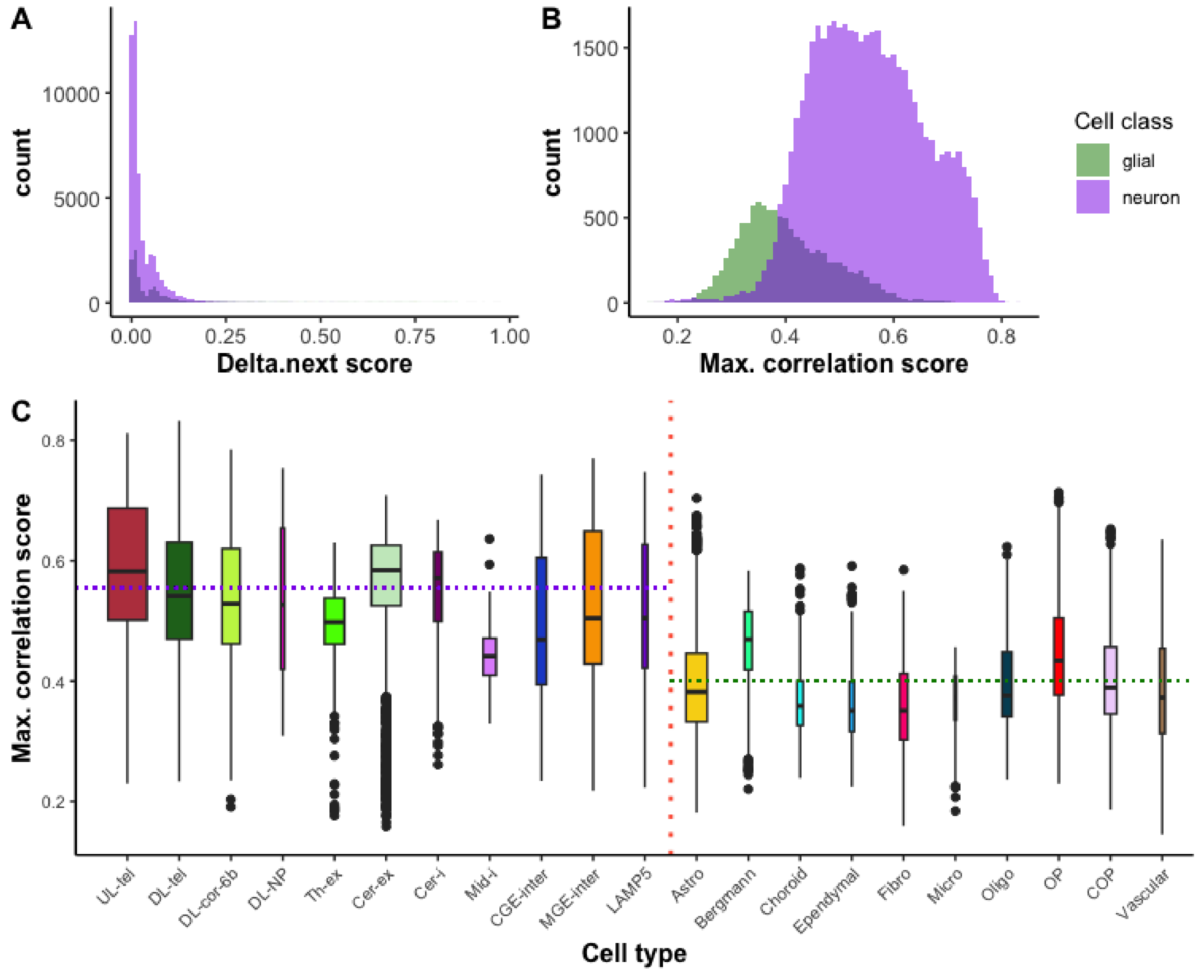


**Figure S2. Cell type annotation using SingleR and annotated human single cell expression data.** (A) Histograms of delta.next scores for glial and neuron cell types, where the delta.next score indicates how much higher the correlation of the best-correlating cell type is compared to the next best when comparing dog and human cell expression. (B) Histograms of maximum correlation scores between dog and human cell expression. (C) Boxplots showing maximum correlation scores between dog and human brain cells, coloured by cell type. Vertical red dotted line splits neurons (left) and glial cells (right). Purple and green horizontal dotted lines indicate mean maximum correlation scores for neuron and glial cell types respectively. Cell type abbreviations are defined in the fig. 1 legend.

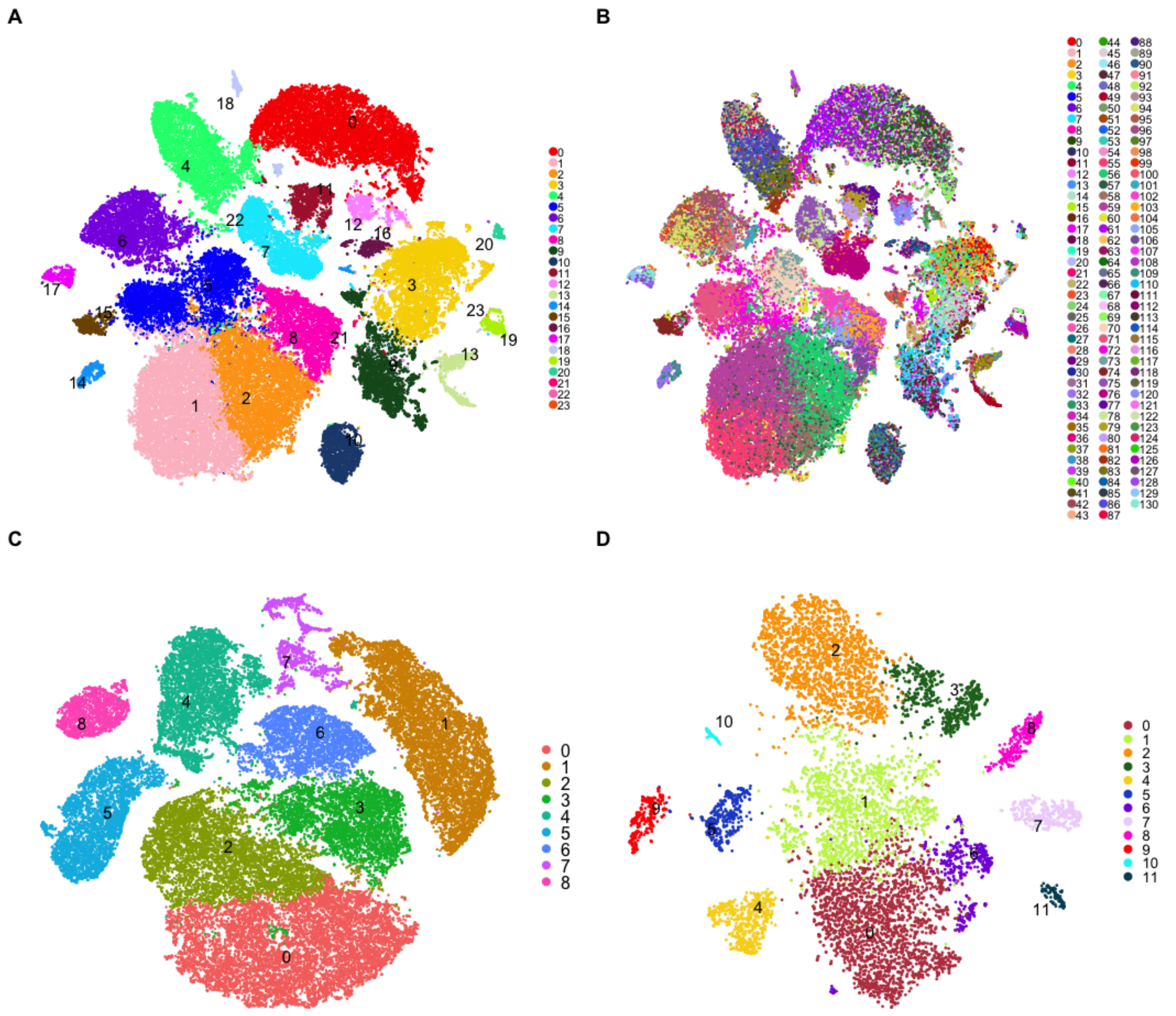


**Figure S3. Clustering of cell types in the dog brain.** Cell clustering was performed using Harmony in Seurat to identify clusters of cells that share similar gene expression profiles. tSNE projections of (A) 24 clusters identified in all cells, (B) 131 subclusters identified among clusters shown in (A), (C) 9 clusters identified among neurons and (D) 12 clusters identified among glial cells.

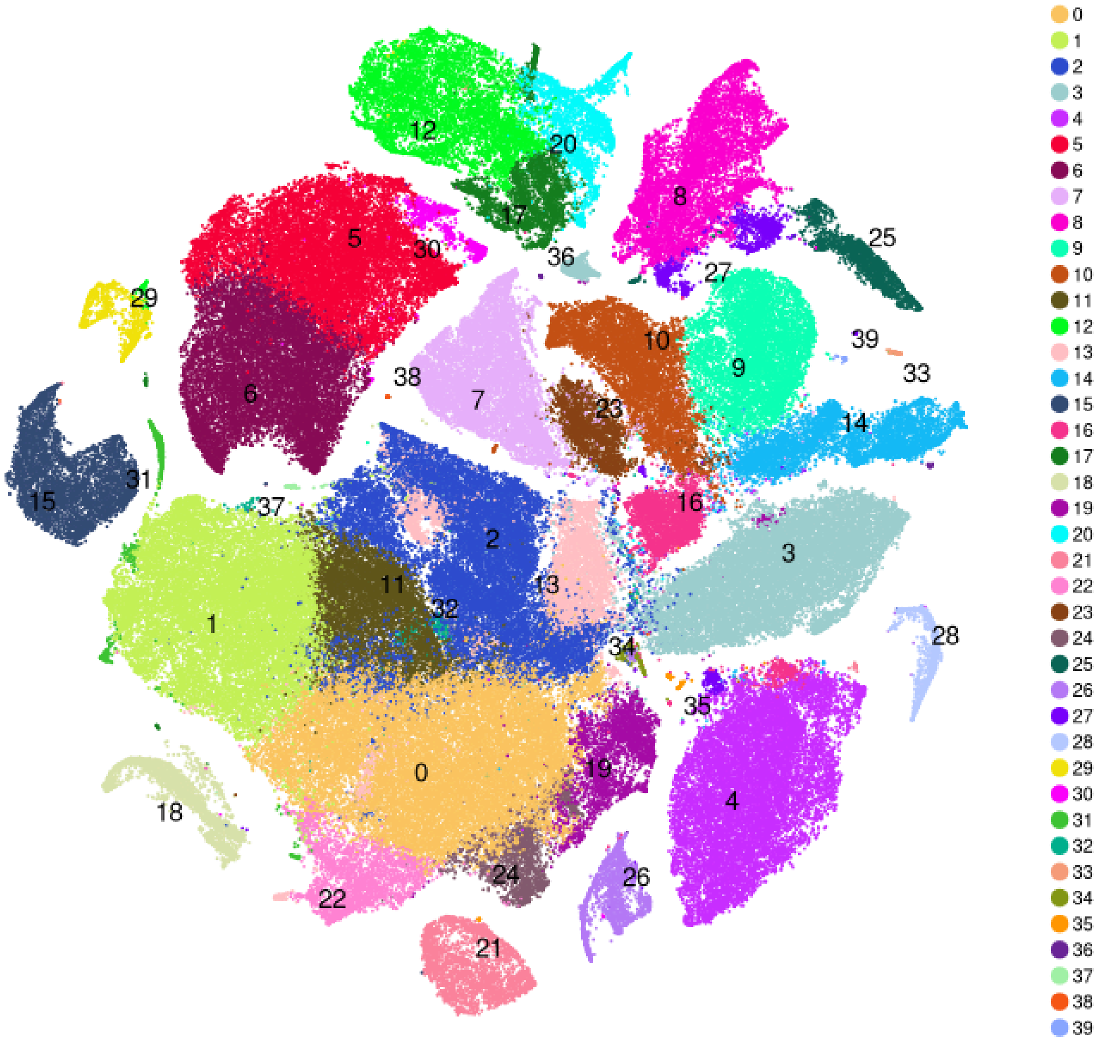


**Figure S4. Clustering of integrated dog-human-mouse data**. 40 distinct cell clusters were identified across the integrated dog-human-mouse data set.

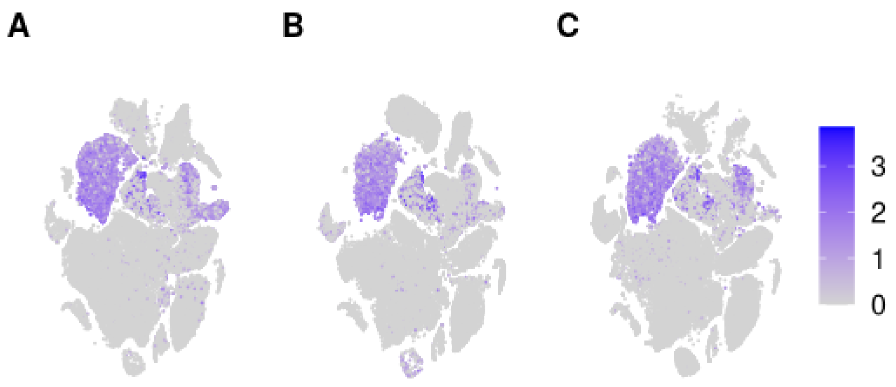


**Figure S5. *RELN* expression in dog, human, and mouse brains.** The gene for the protein reelin shows highly specific expression in cerebellar excitatory neurons in (A) dog, (B) human and (C) mouse brains suggesting a conserved function.

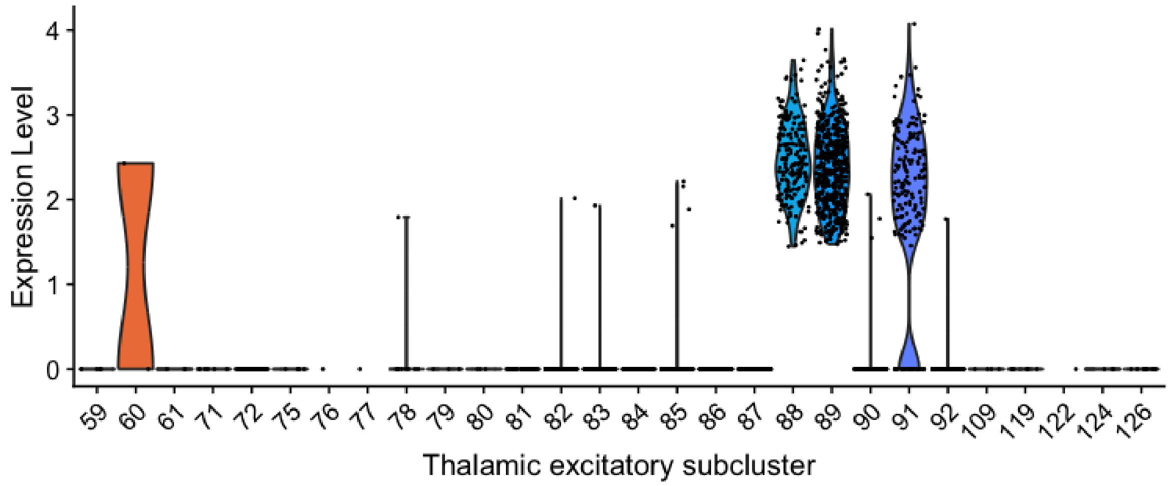


**Figure S6. Expression of *ST8SIA6* in subclusters of thalamic excitatory neurons.** High expression is observed specifically in three subclusters of thalamic excitatory neurons (88,89, and 91). Variation in the second intron of this gene has been associated with the level of sociability towards humans in dogs.

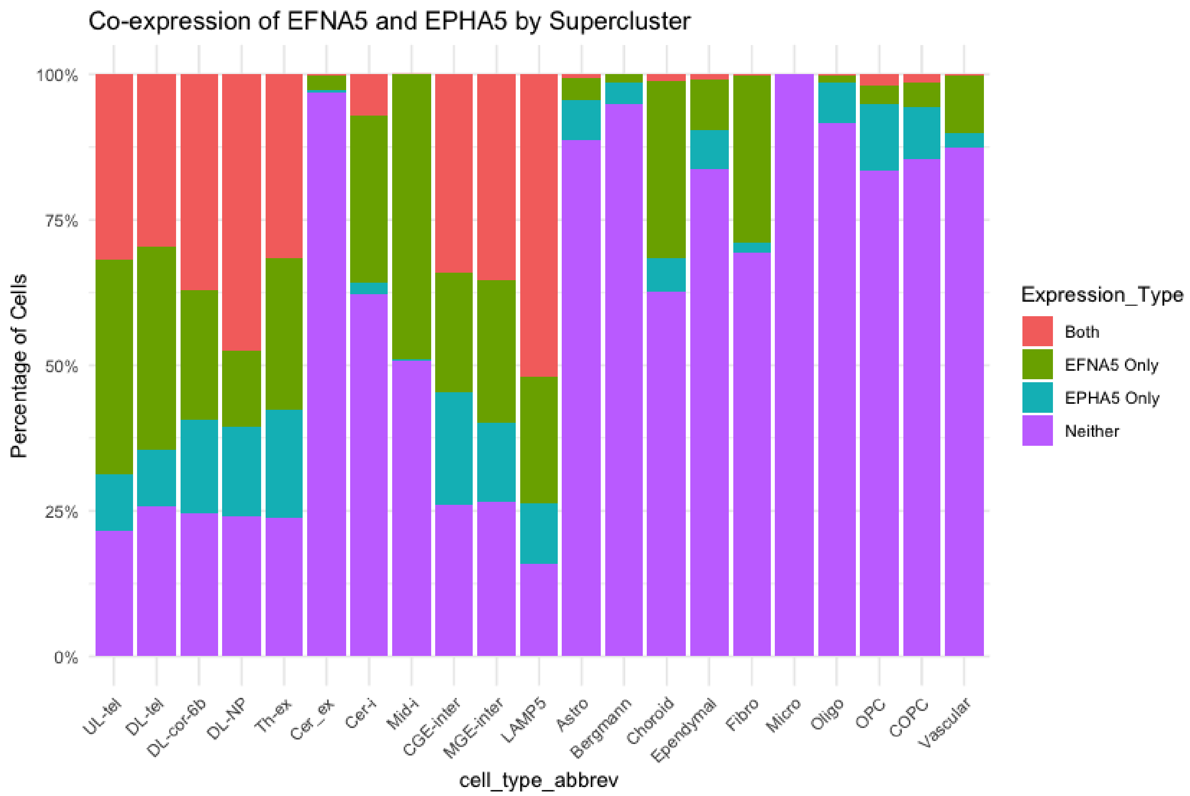


**Figure S7. Coexpression of herding behaviour-related *EFNA5* and *EPHA5* genes in dog brain cells.** We see co-expression of the ephrin ligand gene *EFNA5* and its receptor gene *EPHA5* across all neuron types except for neurons of the cerebellum and midbrain-derived inhibitory neurons. These genes show little or no expression in glial cells. Cell type abbreviations are defined in the fig. 1 legend.
